# Biological and Genetic Determinants of Glycolysis: Phosphofructokinase Isoforms Boost Energy Status of Stored Red Blood Cells and Transfusion Outcomes

**DOI:** 10.1101/2023.09.11.557250

**Authors:** Travis Nemkov, Daniel Stephenson, Eric J. Earley, Gregory R Keele, Ariel Hay, Alicia Key, Zachary Haiman, Christopher Erickson, Monika Dzieciatkowska, Julie A. Reisz, Amy Moore, Mars Stone, Xutao Deng, Steven Kleinman, Steven L Spitalnik, Eldad A Hod, Krystalyn E Hudson, Kirk C Hansen, Bernhard O. Palsson, Gary A Churchill, Nareg Roubinian, Philip J. Norris, Michael P. Busch, James C Zimring, Grier P. Page, Angelo D’Alessandro

## Abstract

Mature red blood cells (RBCs) lack mitochondria, and thus exclusively rely on glycolysis to generate adenosine triphosphate (ATP) during aging in vivo or storage in the blood bank. Here we leveraged 13,029 volunteers from the Recipient Epidemiology and Donor Evaluation Study to identify an association between end-of-storage levels of glycolytic metabolites and donor age, sex, and ancestry-specific genetic polymorphisms in regions encoding phosphofructokinase 1, platelet (detected in mature RBCs), hexokinase 1, ADP-ribosyl cyclase 1 and 2 (CD38/BST1). Gene-metabolite associations were validated in fresh and stored RBCs from 525 Diversity Outbred mice, and via multi-omics characterization of 1,929 samples from 643 human RBC units during storage. ATP and hypoxanthine levels – and the genetic traits linked to them – were associated with hemolysis in vitro and in vivo, both in healthy autologous transfusion recipients and in 5,816 critically ill patients receiving heterologous transfusions, suggesting their potential as markers to improve transfusion outcomes.

**eTOC and Highlights:** 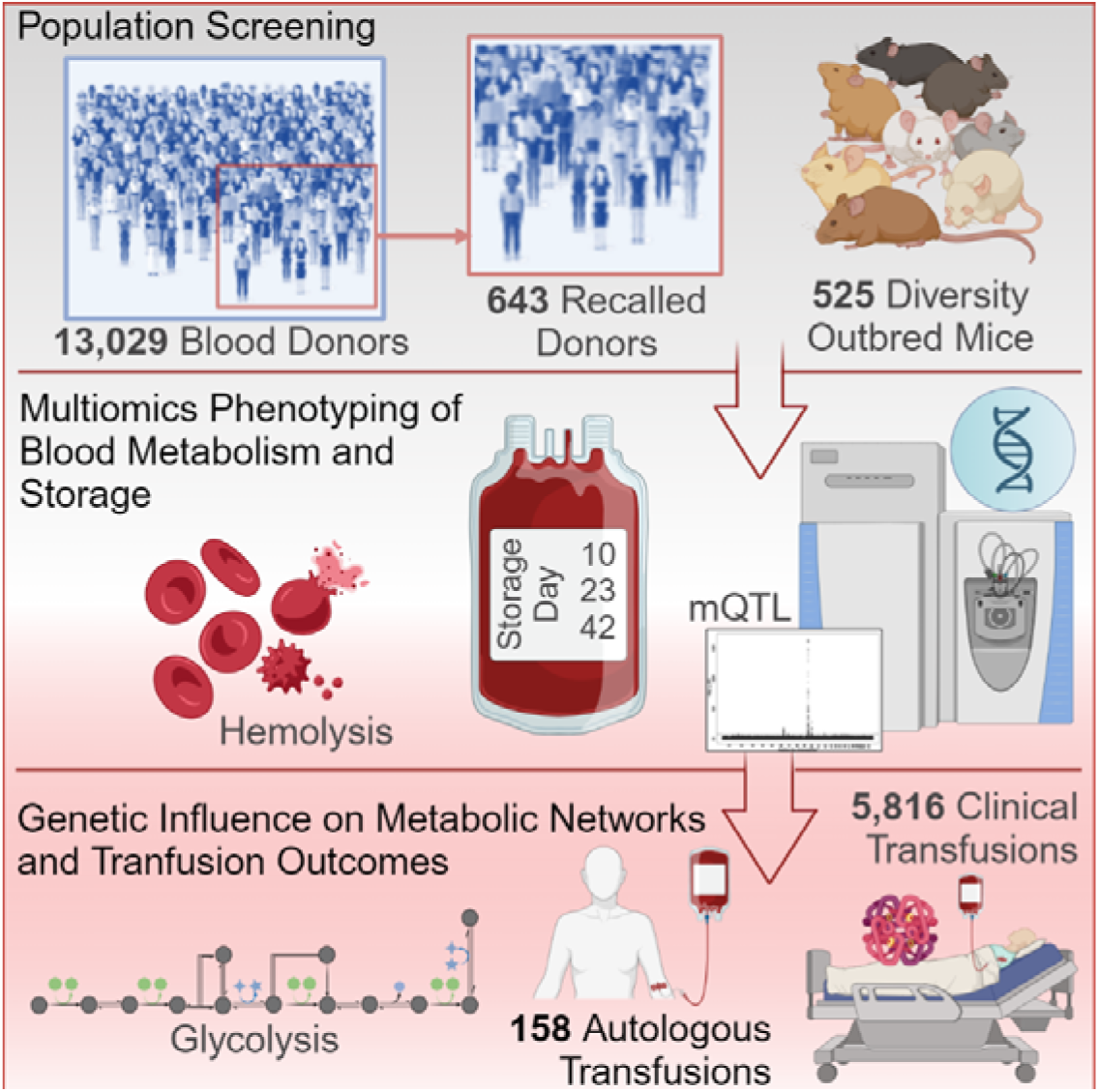

**Highlights:**

- Blood donor age and sex affect glycolysis in stored RBCs from 13,029 volunteers;
- Ancestry, genetic polymorphisms in PFKP, HK1, CD38/BST1 influence RBC glycolysis;
- Modeled PFKP effects relate to preventing loss of the total AXP pool in stored RBCs;
- ATP and hypoxanthine are biomarkers of hemolysis in vitro and in vivo.

## INTRODUCTION

Red blood cells (RBCs) comprise ∼83% of all cells in the human body^1^. Unlike other vertebrates,^2^ mammalian RBCs evolved to maximize oxygen-carrying capacity by increasing hemoglobin content per cell, a feature achieved by discarding nuclei and organelles, including mitochondria. As such, mature RBCs are completely reliant on glycolysis for the generation of adenosine triphosphate (ATP). ATP fuels all key processes in mature RBCs, from proton pumps^3^ to the integrity of membrane proteins^4^ and lipids^5^, from proteasomal activity^6^ to vesiculation.^7^ It has long been posited that ATP-depleted RBCs are rapidly removed from the bloodstream^8^ via mechanisms of intra– or extravascular hemolysis.^3^

Given their lack of *de novo* protein synthesis capacity, RBCs rely on dynamic metabolic reprogramming in response to environmental stimuli, including hypoxia or oxidant stress (e.g., altitude, exercise or storage in the blood bank)^9^. Generation of high-energy phosphate compounds ATP and 2,3-bisphosphoglycerate (BPG) is also central to RBCs’ capacity to perform their main task, i.e., oxygen transport and delivery to tissues. Specifically, ATP and BPG play critical roles in hemoglobin allostery^10^ by stabilizing the tense, deoxygenated state and promoting oxygen release to tissues.^8^

Ease of collection, shear abundance and relative simplicity have made RBCs an attractive eukaryotic cell model for investigating metabolism since the early days of biochemistry.^11^ It is not by coincidence that the earliest efforts to reconstruct human metabolic networks *in silico* have focused on RBC metabolism.^12^ A culmination of nearly a century of studies involving many pioneers in the burgeoning field of biochemistry,^13^ glycolysis – or the Embden-Meyerhof-Parnas pathway – explained the cell’s ability to convert glucose into ATP even in the absence of oxygen.^14^ Despite the importance of metabolism and glycolysis in health and disease, and decades of research on the biochemical determinants of glycolytic fluxes in mammals,^15^ understanding glycolytic regulation in mature RBCs holds relevant implications for systems physiology (e.g., gas exchange), and clinical practice. Indeed, understanding erythrocyte metabolism holds the key to optimizing storage conditions and predicting post-transfusion performance of blood products.^16^

Transfusion of packed RBCs (pRBCs), the most common in-hospital medical procedure after vaccination, is a life-saving intervention for tens of millions of civilian and military patients whose survival depends solely on chronic or massive transfusions, with no current alternative medical interventions. Altered energy metabolism is a hallmark of the so-called storage lesion,^17^ a series of biochemical and morphological alterations that ultimately increase the propensity of stored RBCs to hemolyze *ex vivo* in the refrigerated unit or *in vivo* upon transfusion. Biopreservation of pRBCs for up to 42 days at 4°C poses an energetic challenge to erythrocytes, as proton pumps fail under refrigerated storage temperatures and become a sink for ATP consumption.^17^ Storage-dependent consumption of glucose is not limited by its availability in currently licensed storage additives, which contain supra-physiological concentrations of glucose in all the most common additives (e.g., 111 mM and 55 mM for additive solutions 1 and 3 in the United States – AS1 and AS3, respectively).^18^ As storage progresses and glucose is metabolized through glycolysis, lactate accumulates up to 35 mM in the closed system of a blood bag,^19^ with a concomitant drop in pH.^20^ Intracellular acidification has a negative inhibitory feedback on pH-dependent enzymes, including phosphofructokinase (PFK), BPG mutase (BPGM), and glucose 6-phosphate dehydrogenase (G6PD), the rate-limiting enzymes of glycolysis, the Rapoport-Luebering shunt and the pentose phosphate pathway (PPP), respectively.^21^ Combined with slower enzyme kinetics at 4°C (compared to 37°C *in vivo*),^22^ ATP synthesis rates become insufficient to sustain fueling of proton pumps. As a result, BPG pools (∼5 mM) rapidly deplete to promote compensatory ATP synthesis, with BPGM phosphatase activity favored over its kinase one by the drop in pH.^23^ As adenylate pools shift to lower energy adenosine monophosphate (AMP) and adenosine, BPG-depletion and oxidant stress both contribute to activate adenosine/AMP deaminase 3,^24,25^ which ultimately generates hypoxanthine,^22,24,26,27^ a key hallmark of the metabolic storage lesion.^17^ Although the clinical relevance of the storage lesion remains unresolved – which is, in part, the focus of this study – it is now clear that RBC metabolism under refrigerated storage conditions is impacted by storage temperatures, additive solutions, donor dietary or other exposures (e.g., smoking, alcohol consumption, drugs that are not grounds for blood donor deferral), and also by factors such as donor age, sex, ethnicity and body mass index (BMI).^22,24,26–28^ While it has been posited that RBC storability and post-transfusion performance are a function of donor genetics^29^ and non-genetic factors,^30^ the biological and genetic underpinnings of RBC glycolysis have not yet been determined, since no study to date was sufficiently powered to test the independent and combined contributions of these factors from an omics standpoint, which is the central focus of this study.

Current gold standards of storage quality are the measurement of (i) “in bag” hemolysis (i.e., the percentage of RBCs that lyse spontaneously by storage in the blood bank for 42 days) and (ii) post-transfusion recovery (PTR) determined by radiolabeled ^51^Cr studies in healthy recipients, which measures the percentage of stored RBCs still circulating in the bloodstream of the recipient 24h after autologous transfusion. The FDA and European Council require <1% and >75% thresholds for end-of-storage hemolysis and PTR, respectively, as the gold standards of quality for the last several decades. However, these are limited to licensing new products, as these parameters are not routinely tested on all products issued for transfusion, owing to logistical and economic considerations. In addition, both metrics have major limitations,^31^ and inter-donor heterogeneity in these parameters has been noted for both.^32–34^ Finally, neither of these “quality” testing parameters provides insights regarding the preservation of RBC “function” upon storage, which is a critical unmet clinical need. For these reasons, both the National Heart, Lung and Blood Institutes^35^ and the Food and Drug Administration (FDA)^36^ have listed the identification of new functional biomarkers of RBC storage quality and post-transfusion performances as central to their programmatic agenda.

Here we combined ultra-high-throughput metabolomics (1 min per sample) and genomics approaches to test the hypothesis that energy metabolism can be monitored as a biomarker of RBC storage quality. Specifically, we hypothesize that markers of ATP synthesis, consumption and deamination inform on RBC hemolytic propensity, either “in the bag”, or in the circulatory system of the recipient following transfusion. We further test the role of donor biology, including age, sex and genetic polymorphisms in regulating stored RBC metabolism. In summary, the present study addresses knowledge gaps that remain in understanding (i) how biological and genetic factors impact stored RBC metabolism; (ii) whether RBC metabolism is associated with transfusion outcomes; and (iii) whether metabolites in this pathway provide useful biomarkers of RBC storage quality and transfusion efficacy.

## RESULTS

### Glycolysis is influenced by donor sex, age, ethnicity and BMI

First, we performed high-throughput metabolomics analyses on leukocyte-filtered pRBCs, which were donated by 13,029 volunteers at four different blood centers across the United States and stored for 42 days (i.e., at the end of the FDA-approved shelf-life – **Figure 1.A**). The donor population was balanced with respect to sex (6,507 males and 6,522 females) and blood center (from a minimum of 3,019 to a maximum of 3,564 per center). The age range spanned from 18 to 87 (median age 47) with median body mass index (BMI) of 26.6. Most donors were of Caucasian descent (7,037), with substantial representation of individuals of Asian (1,602), African (1,542) and Hispanic descent (1,153) and “Other”^29^ ethnicities (including multiracial people, Alaska/Hawaiian/Native Americans, and Pacific Islanders – 1,695). RBCs were exclusively stored in AS-1 and AS-3 (compositions detailed in **Data S1**). Linear discriminant analysis (LDA) as a function of donor age, sex, BMI and ethnicity suggested their impact on the metabolic heterogeneity of end-of-storage pRBCs (**Figure 1.B-E**).

**Figure 1.**
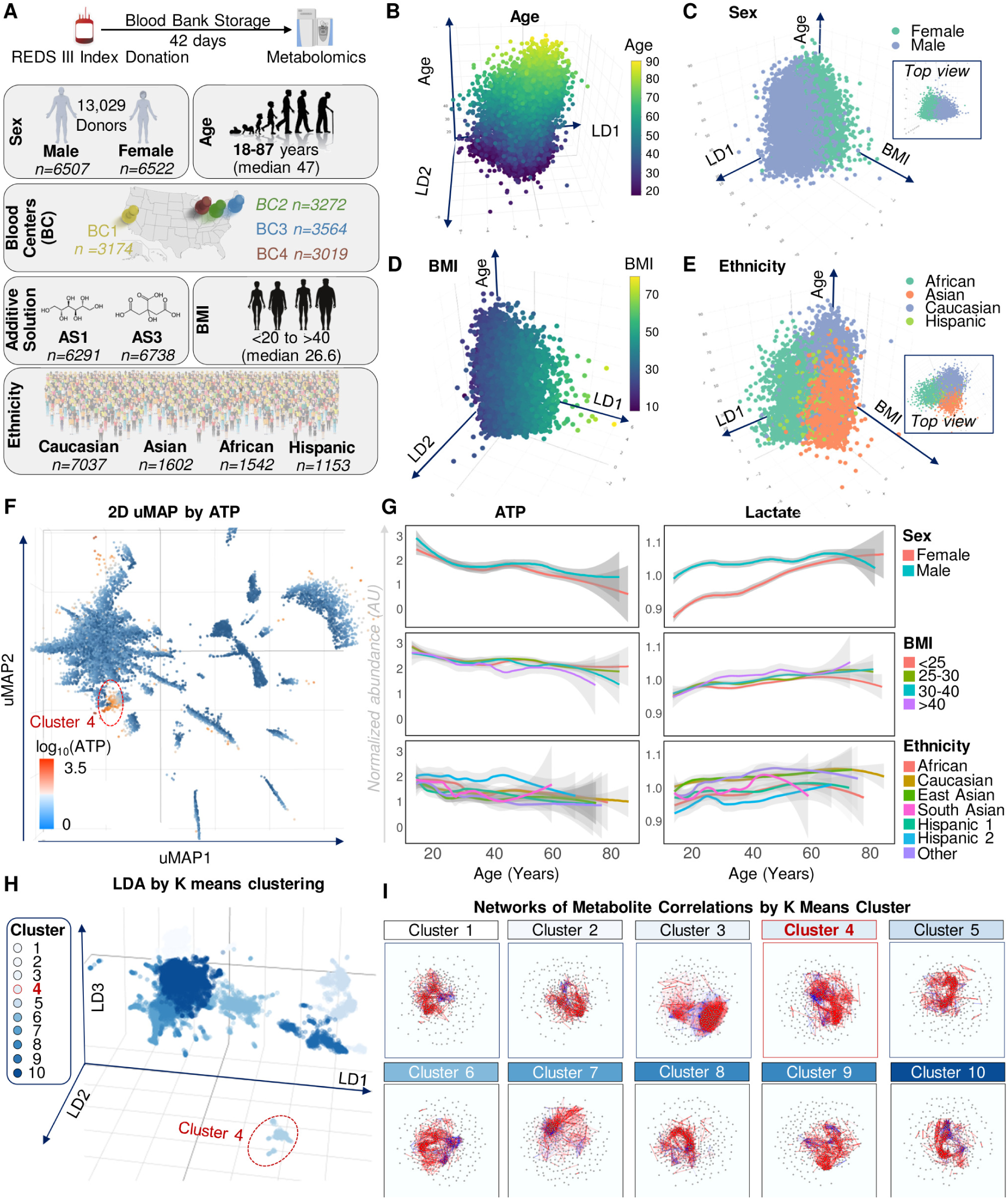
– Metabolomics of 13,029 end of storage packed RBCs from the REDS RBC Omics Index cohort. Overview of the study design and donor demographics (sex, age, ethnicity) (**A**). In **B-E**, linear discriminant analysis (LDA) of metabolomics data a function of donor age, sex, BMI and ethnicity. In **F**, uMAP identifies a cluster of donors with elevated ATP. In **G,** donor age, sex and ethnicity, but not BMI, significantly impact the levels of ATP and lactate in end of storage RBCs. In **H**, K-means clustering analysis highlights separation of cluster 4 among the top 10 clusters. In **I**, network view of metabolite-metabolite correlations (Spearman) across clusters. Each metabolite is a node; red/blue edges indicate positive/negative correlations >|0.5|.

Unsupervised uniform manifold approximation and projection (uMAP) was used to cluster the 13,029 samples based on metabolic phenotypes alone, which highlighted a group of donors with elevated ATP (**Figure 1.F**). Younger donors had significantly higher levels of ATP, which was negatively associated with donor age, independently of sex, with lowest levels observed in donors severe obesity (BMI>40); higher levels of ATP were instead observed in donors of Hispanic descent (Mexican and Central American Hispanics), followed by Caucasians, even at older ages (**Figure 1.G**). Lactate was higher in older, male donors of Caucasian, East Asian or Other descent. Negative associations were observed between age and the levels of ATP and all glycolytic intermediates, except lactate (**Supplementary Figure 1.A**). Despite consistent age-associated trends for all donors, sex dimorphism was observed, with lower levels of glucose and higher levels of glycolytic metabolites in males across the age range; sex dimorphism was lost for donors past 50 years of age, the average age of menopause in the United States^37^ (**Supplementary Figure 1.B**). RBCs from donors with the highest BMIs (>40) were characterized by significantly higher levels of lactate, as compared to all the other BMI groups within the 30-60 age window (**Supplementary Figure 1.C**).

### Unsupervised clustering separates donors with high glycolysis from the rest of the donor population

K-means clustering separated donors based on metabolic patterns of end of storage units, especially cluster 4, the most distinct amongst all clusters (**Figure 1.H**). Network analyses of metabolite-metabolite correlations were performed for each cluster separately (**Figure 1.I**) or combined (**Figure 2.A**), showing the centrality of glycolytic metabolites and their association to donor demographics, especially age and sex (**Figure 2.A**). Hierarchical clustering analysis of ANOVA across clusters confirmed a strong signal separating donors in cluster 4 from the rest (highlighted in red in **Figure 2.B**). Donors in cluster 4 were characterized by the highest levels of ATP and total adenylate pools (AXP – calculated as (ATP+0.5*ADP)/(ATP+ADP+AMP) through the whole donor age range (**Figure 2.C; Data S1**), suggesting that higher ATP levels in this group are not exclusively explained by donor age. Cluster 4 donors also showed the highest levels of all glycolytic metabolites (**Figure 2.D**) and lowest levels of hypoxanthine (HYPX), the top positive and negative correlates to ATP levels (**Supplementary Figure 2**). Donors in clusters 4 were overall younger, leaner (lower BMI), with a highly proportion of male donors compared to the other clusters combined (**Figure 2.E**). These units were mostly stored in AS-3 (87%) versus AS-1 (13%), while the rest of the population had a comparable balance (51 vs 49%) between the two additives (**Figure 2.E**). Consistent with the literature^19,22,26,27,38–42^ and the composition of the additives (violin plots in **Data S1**), higher glucose levels in AS-3 were associated with higher levels of ATP, BPG and all glycolytic metabolites for donors of all ages, with the exception of lactate (**Figure 2.F**). This observation differs from the analyses based on donor demographics, in which ATP, glycolysis and lactate levels were concordant (all higher in males than females – **Supplementary Figure 1.B**). Additives alone are insufficient to explain the boost in glycolysis, since 360 units in cluster 4 were stored in AS-3 (80%), against a residual 6,378 from all the other groups combined (**Data S1**). Donors in cluster 4 had lower storage hemolysis and were significantly enriched in donors of Caucasian or Hispanic descent, with significantly lower representation of donors from other ancestries (**Figure 2.E**)

**Figure 2.**
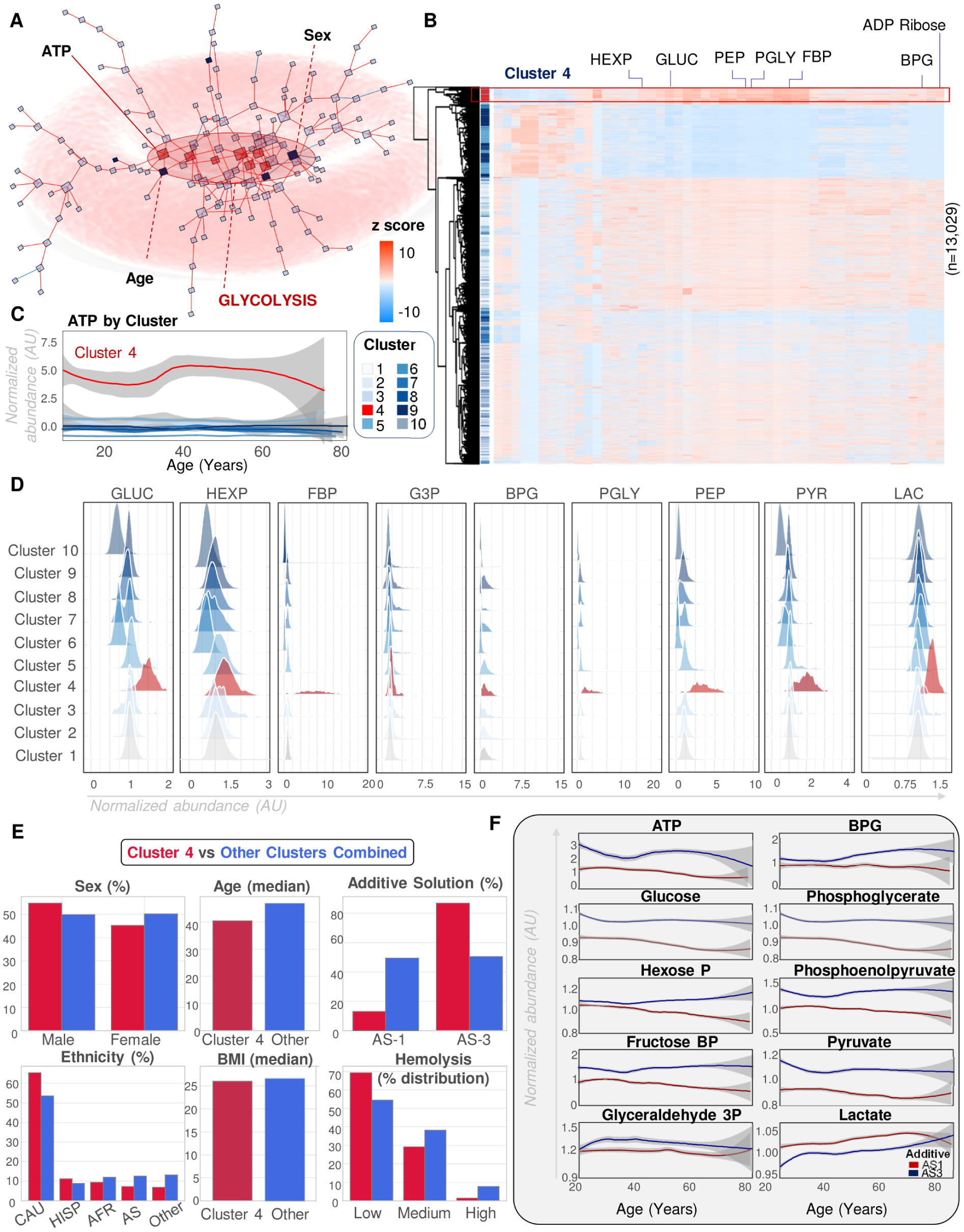
– Cluster 4 donors have elevated glycolysis, ATP and low hypoxanthine. Network view. (**A**) of the most significant metabolite-metabolite interactions across the whole dataset confirms the centrality of glycolytic metabolism in stored RBCs. In **B**, hierarchical clustering analysis (top 50 metabolites by ANOVA) of index donors based on K-means clustering (cluster 4 in red). Glycolytic metabolites are abbreviated, including glucose (GLUC), hexose phosphate (HEXP), fructose bisphosphate (FBP), glyceraldehyde 3-phosphate (G3P), 2,3-bisphosphoglycerate (BPG), phosphoglycerate (PGLY), phosphoenolpyruvate (PEP), pyruvate (PYR), lactate (LAC). In **D**, ridge plots of glycolytic metabolites (highest in cluster 4 – red). In **E**, donor demographics and hemolysis for cluster 4 (red) vs all the other clusters combined (blue). In **F**, glycolysis by storage additive (AS1 in red, AS3 in blue).

By study design, the REDS RBC Omics project included a validation phase in which 643 donors (out of 13,029), who ranked in the 5^th^ or 95^th^ percentile for end-of-storage measurements of hemolysis at the index phase, were invited to donate a second RBC unit^33^ for multi-omics analyses at storage days 10, 23 and 42.^43^ The steady state abundance of glycolytic metabolites at storage day 42 was significantly reproducible across both donations from the same (643) donors at the index and recalled phase (**Supplementary Figure 2.F; Data S1**), suggesting a potential impact of genetics, in addition to biological factors.

### Genetic underpinnings of glycolytic heterogeneity in fresh and stored murine RBCs in the J:DO mice

To delve into the genetic underpinnings of metabolic changes while controlling for biological (sex, age, BMI) and processing factors (storage duration and additives), we leveraged the Jackson Laboratory’s Diversity Outbred (J:DO) mouse population. Briefly, eight genetically diverse founder strains were crossed for >46 generations to establish this population with highly recombined genomes (**Figure 3.A**). These 525 mice were genotyped on an array with 137,192 markers; fresh and stored RBCs were tested for levels of glycolytic metabolites, and results were adjusted for sex and cage batches (**Figure 3.A**). Murine RBCs were stored for 7 days, which was chosen following pilot studies aimed at identifying a storage range that resulted in end-of-storage PTR comparable to humans^44^, while also accounting for the wide variability in PTR amongst the J:DO founder mouse strains. Our analyses identified key regions of co-mapping metabolite Quantitative Trait Loci (mQTLs – **Data S1**) on murine chromosomes 5 and 13 for ATP levels and glycolytic metabolites, highlighted in Manhattan plot, hive plot and heat map illustrations (**Figure 3.B-D**). Representative Manhattan plots, founder haplotype effects and locus zoom plots are shown for ATP and fructose bisphosphate (FBP) in **Figure 3.E-H**. Altogether, these analyses associate ATP and glycolytic metabolites with a polymorphic region on chromosome 5 that encodes for the ectonucleotidase CD38, also known as ADP-ribosyl cyclase 1, or the neighboring bone marrow stromal antigen – BST1 (also known as CD157 or ADP-ribosyl cyclase 2), a pH-dependent NAD(P)^+^ hydrolase.^45,46^ Glycolytic metabolites and ATP were also associated with a second region on murine chromosome 13, which encodes phosphofructokinase platelet – PFKP (**Figure 3.G-H**). However, while the CD38/BST1 region was associated with all glycolytic and PPP metabolites in stored and fresh murine RBCs, the PFKP region was only linked to stored RBC metabolites (**Figure 3.B; Supplementary Figure 3.A-D**).

**Figure 3.**
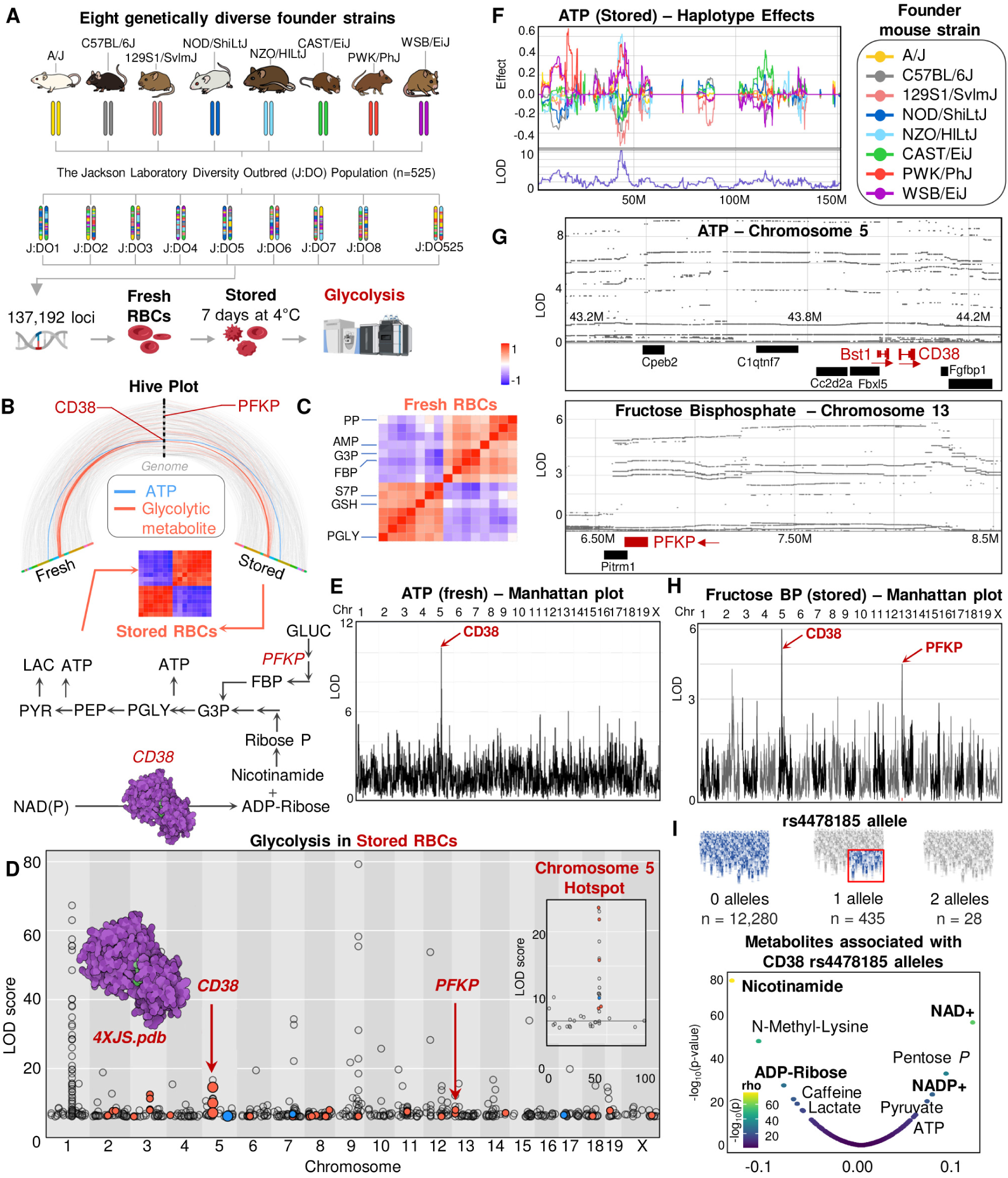
– Glycolysis and mQTL in fresh and stored mouse RBCs from the Diversity Outbred population. Metabolomics analysis was performed in fresh and stored RBCs from 525 mice derived from crosses of 8 genetically-distinct founder strains (Jackson Laboratory’s Diversity Outbred – J:DO – **A**). Hot spots of mQTLs were identified on chromosomes 5 and 13 – hive plot (**B**). Heat maps of metabolic correlates to the locus on chromosome 5 in fresh (**C**) and stored RBCs (**B**). Manhattan plot with highlighted a hotspot region on chromosome 5 (**D**), which codes for CD38 and associates with all glycolytic and pentose phosphate pathway (PPP) metabolites in stored murine RBCs. CD38/BST1 was associated with ATP levels in fresh and stored murine RBCs (Manhattan plot, Best Linear Unbiased Predictors and Locus Zoom in **E-F-G,** respectively). Both CD38 and PFKP were associated with fructose bisphosphate (FBP) levels in stored RBCs (locus Zoom in **G,** and Manhattan plot in **H**). In **I,** association between the CD38 SNP rs4478185 and NAD(P)^+^, glycolysis/PPP metabolites in human RBCs from the REDS RBC Omics Index cohort.

Haplotype effects in the BST1/CD38 region reveal an impact of alleles inherited from NZO/ HlLtJ, PWK/ PhJ, and WSB/Eij strains (e.g., for ATP – **Figure 3.F**), which are inverted for Glyceraldehyde 3P in stored RBCs (**Supplementary Figure 3.E-G**). Haplotype effects for the mQTL in the PFKP region for FBP in stored RBCs are more complex and distinct from the CD38 region (**Supplementary Figure 3.B**).

In the REDS RBC Omics Index cohort, the CD38 SNP rs4478185 – while rare (2 and 1 alleles in 0.2% and 3.4% of the index population, respectively) – was significantly associated with levels of glycolytic metabolites and ATP (**Figure 3.I**). As an internal validation of the quality of the metabolomics data, we observed a positive association of this SNP with NAD(P)^+^, and a negative association with nicotinamide and ADP-ribose, substrates and products, respectively of the enzymatic activity of CD38 (**Figure 3.I**). Indeed, mQTL analysis for ADP-ribose highlighted both CD38/BST1 and PFKP SNPs as top loci associated with the end of storage levels of this metabolite in the index donor population (**Figure 3.I**).

### Genetic polymorphisms associated with the glycolytic heterogeneity in 13,029 REDS donors

To help further translate these murine findings into the human settings, we tested the 13,029 REDS Index donors for ∼879,000 single nucleotide polymorphisms (SNPs), which enabled mapping of mQTL for glycolytic metabolites (**Figure 4.A, Data S1**), upon adjustment for donor sex, age, BMI and storage additives. In the GWAS catalogue^47^ previously reported SNPs have been linked to hematological parameters, such as RBC count, hematocrit, platelet (PLT) count, and hemoglobin concentration (**Data S1**). A summary Manhattan plot of mQTL results is shown for all glycolytic metabolites merged together (**Figure 4.B**), or just for ATP (**Supplementary Figure 4**). Manhattan plots are also shown for glycolytic metabolites and hypoxanthine (**Figure 4.C-J**). A similar analysis was performed for the PPP end product, pentose phosphate (including ribose 5-phosphate and other pentose phosphate isomers that are not resolved by the high-throughput assay). **Supplementary Figure 5** shows the locus zoom plots for the most common PFKP SNPs (chromosome 10) that associate with ATP, hypoxanthine and lactate (FDR-corrected p-value = e-163).

**Figure 4.**
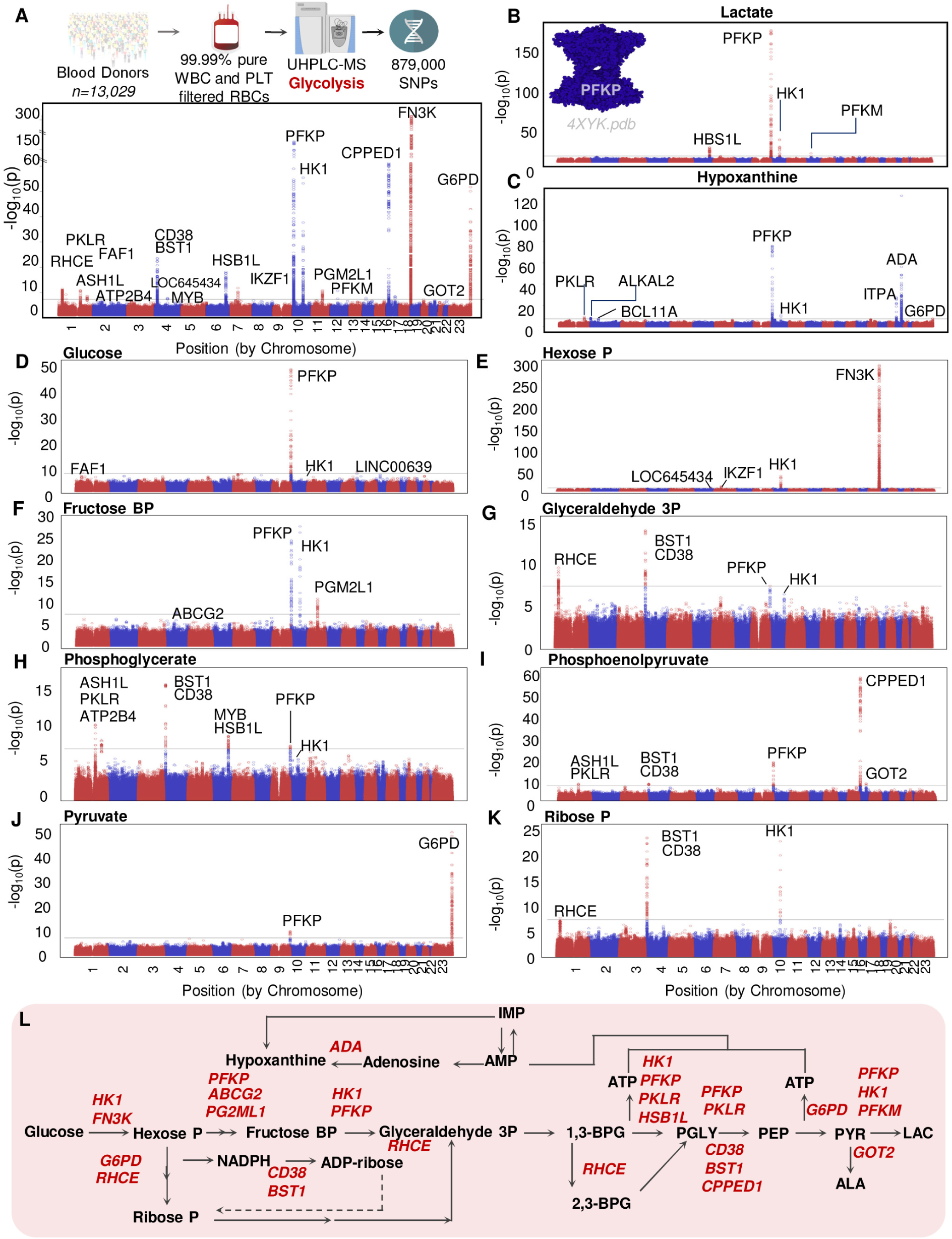
– Genetic underpinnings of glycolytic heterogeneity in stored RBC units from 13,029 donors. Metabolite quantitative trait loci (mQTL) analyses were performed for all glycolytic metabolites. Results are shown as Manhattan plots for combined results (**A**), or for separate metabolite (lactate (**B**), hypoxanthine (**C**), glycolysis and ribose phosphate (**D-K**). Abbreviations: P: phosphate; BP: bisphosphate; Gene names are listed with their official gene symbols. In **L**, a summary overview of the gene-metabolite associations (official gene symbols are listed in red next to the metabolite they have been linked to).

This analysis also identified multiple common hot spot areas in regions coding for other rate-limiting enzymes of glycolysis,^15^ hexokinase 1 (HK1, chromosome 10), phosphofructokinase M (PKFM, chromosome 12), pyruvate kinase (PKLR, chromosome 1), or other enzymes involved in glucose 1,6-bisphosphate metabolism (phosphoglucomutase 2 like 1 – PGM2L1, chromosome 11 – **Figure 4.C** and **I**). Similarly, end of storage levels of pyruvate showed a strong association with the X-linked rate-limiting enzyme of the PPP, glucose 6-phosphate dehydrogenase (G6PD –**Figure 4.D**). For glycolytic metabolites downstream to glyceraldehyde 3-phosphate, a common hit – absent in the hexose phosphate group – was CD38/BST1. Both CD38 and BST1 map to human chromosome 4, and multiple distinct loci in this region were linked to various metabolites including ADP-ribose, pentose phosphate and NADP^+^ (**Supplementary Figure 5.B-E**). A summary overview of the gene-metabolite associations is shown in **Figure 4.L;** detailed results are in **Data S1.**

### Glycolytic heterogeneity and donor genetic ancestry

To expand on the mQTL analyses, we determined minor allele frequency and direction effects for different genetic ancestry groups (**Data S1**). Despite Caucasian donors being over-represented (54% of the population), both consistent and heterogeneous effects across ancestries were observed. For example, the ATP-PFKP rs115363550 association (**Figure 5.A**) was strongest in Caucasian donors, and almost completely lost significance and disappeared in donors of African descent, while it was not observed in donors of Hispanic and Asian ancestry (**Figure 5.B**). Discordant direction effects were observed between donors carrying two alleles of the rs115363550 SNP versus the rs2388595 (**Data S1**), the latter associated to lactate levels across all genetic ancestries (**Figure 5.C-D**). Some associations (e.g., hexose phosphate and FN3K) were highly significant with consistent effects across all donor ancestries (**Figure 5.E-F**), with strongest signals in donors of Caucasian and Asian descent, followed by Hispanic and African donors. On the other hand, some gene-metabolite associations were unique to specific groups. For example, the association between pyruvate and G6PD (rs1050828, c.202G>A; p.Val68Met), known as the “common African variant”,^48^ was observed exclusively in donors of African descent and, to a lesser extent, in Asian donors (**Figure 5.G-I**).

**Figure 5.**
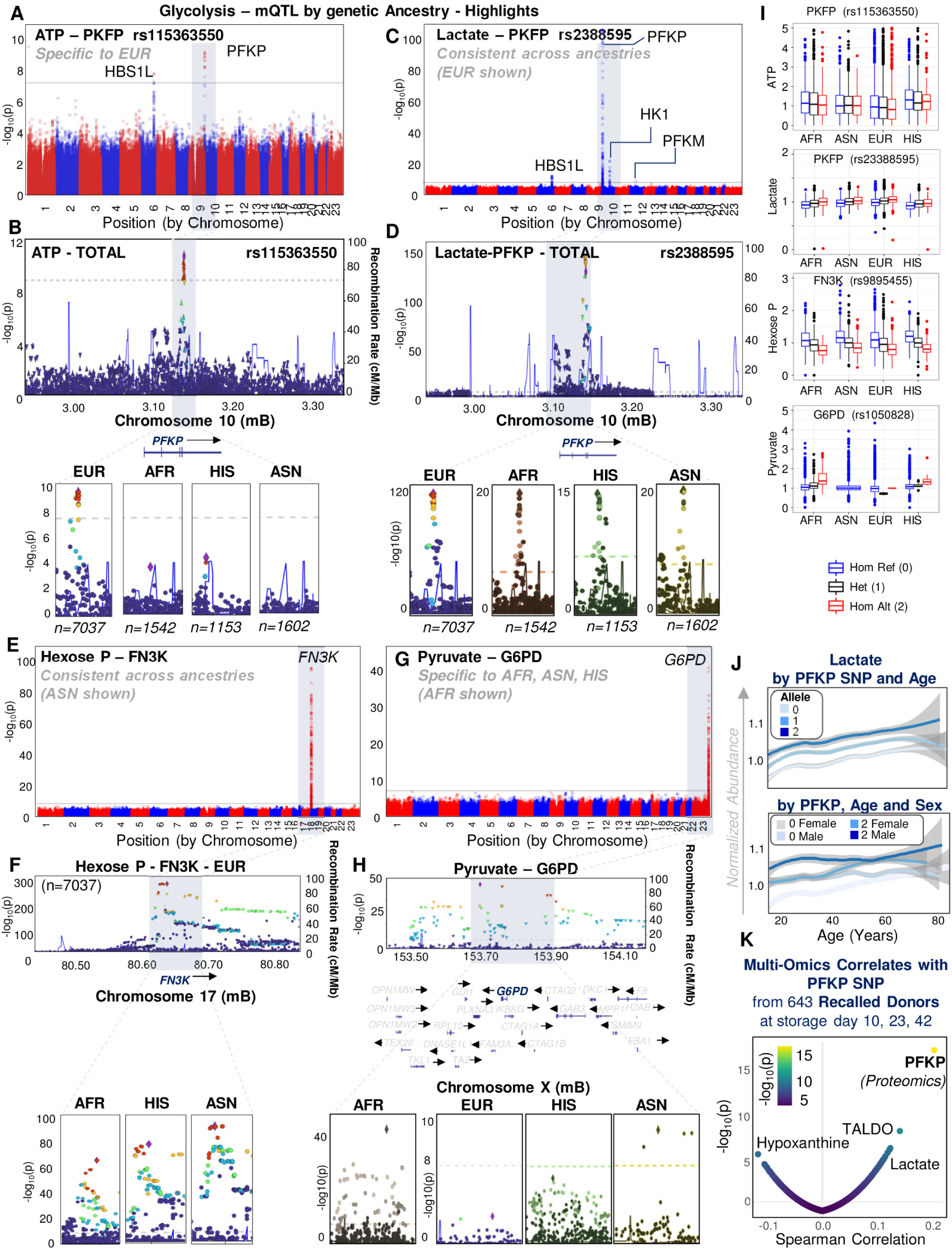
– mQTL analyses by ancestry. In **A-B**, Manhattan plot and locus zoom for genome-wide associations for ATP mapping on PFKP rs115363550, exclusively observed in donors of European descent (EUR) (**B –** bottom panel – AFR: African American; HISP: Hispanic – direction effects in **Data S1**). In **C-D**, Lactate associates to PKFP SNP rs2388595 across ancestries. In **E-F**, hexose phosphate association to multiple FN3K SNPs were consistently observed across ancestries. Conversely, pyruvate association to G6PD SNPs was only observed in donors of African and Asian descent (**G-H**). Summary box plots by ancestries in **I**. In **J**, combined effects of PFKP SNP (0, 1 or 2 alleles) with donor age (top) and sex (bottom) for lactate. In **K**, multi-omics analyses of recalled donor units (n=643, storage day 10, 23, 42) confirmed a strong positive association between PFKP rs2388595 alleles and PFKP protein levels or lactate, and a negative association with hypoxanthine.

### PFKP polymorphisms and donor biology in index and recalled donors

In index donors, the presence of the alternative allele for the top PFKP SNP (rs2388595) was significantly associated with lactate (positive) and hypoxanthine levels (negative) (**Figure 5.J**), and impacted all glycolytic metabolites (**Supplementary Figure 6.B**). The frequency of PFKP rs2388595 allele in homozygosity was highest in donors of Caucasian (25%) or Asian descent (23%), and lowest in donors of Hispanic descent (18% – **Data S1**). A breakdown of minor allele frequency across ancestries for this SNP is summarized in **Supplementary Figure 4.F** (full list for all SNPs in **Data S1**).

To validate and expand on these genomics and metabolomics findings, we performed proteomics analyses on 1,929 samples at storage day 10, 23 and 42 from longitudinal sampling of a second set of 643 pRBC units from the recalled donor cohort (**Data S1**). These analyses showed that PFKP protein levels were the most significant variable positively associated to PFKP rs2388595 allele dosage, suggesting that this intronic variant impacts protein expression (**Figure 5.K**). While explaining a relatively minor percentage of the ATP variance compared to storage additives (0.3% vs 27.7%), the PFKP rs2388595 SNP explained 2.7% and 4.2% of hypoxanthine and lactate variance across all units (**Figure 6.A**). Even though no significant interaction effects were observed between PFKP and biological factors (**Figure 6.A; Supplementary Figure 4.G**), the influence of biological variables – such as age and sex – further impacted metabolite levels in the context of PFKP SNPs, especially with respect to glucose and fructose bisphosphate consumption and lactate accumulation, with concomitantly lower levels of hypoxanthine in older male donors carrying two alleles of PFKP SNP rs2388595 (**Supplementary Figure 6.D-E**). On the other hand, the abundance of PFK protein isoforms was not impacted by donor age and sex (**Supplementary Figure 6.F**).

**Figure 6.**
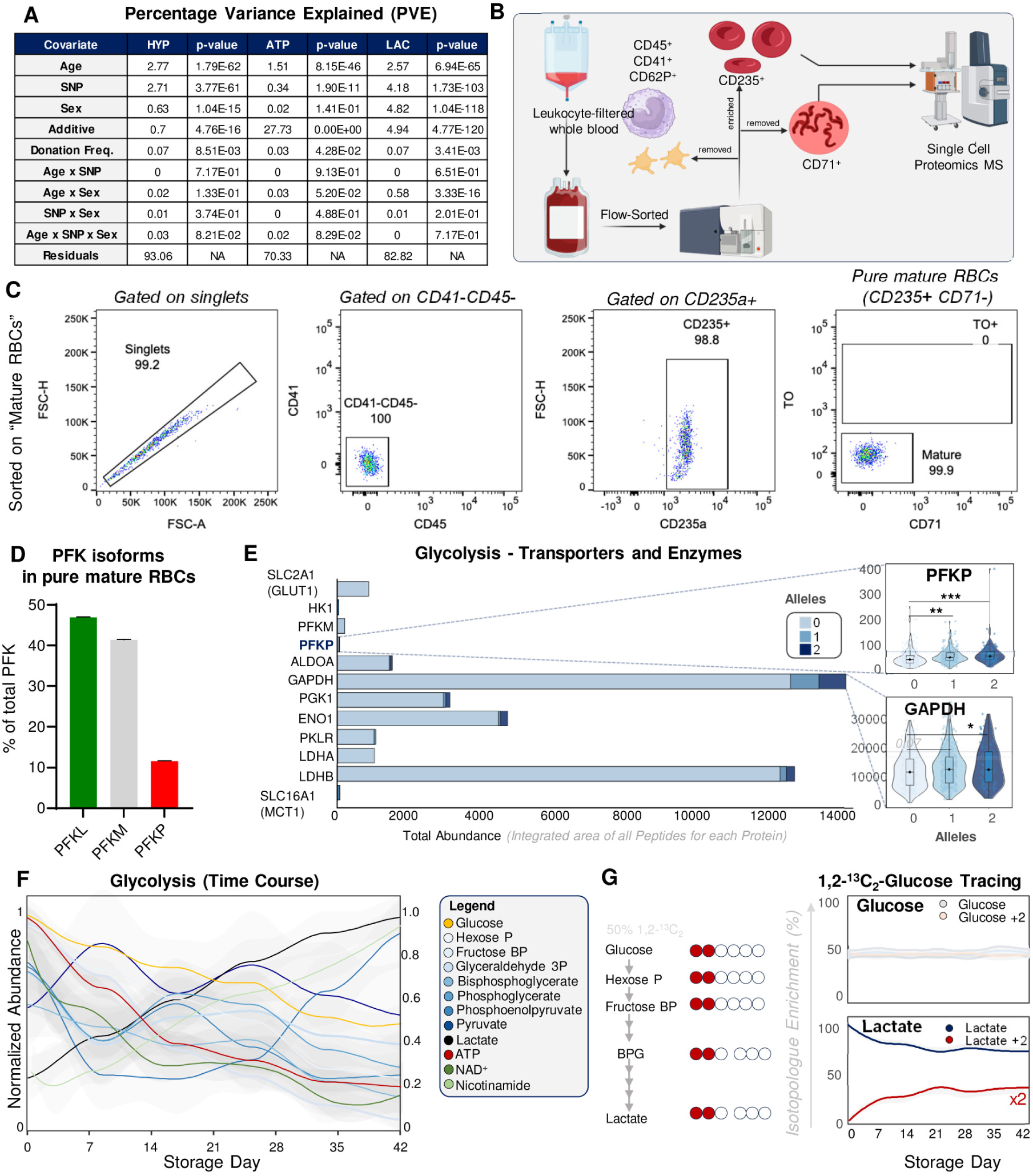
– PFKP is expressed in ultra-pure mature RBCs. Percentage variance explained (PVE) for ATP, hypoxanthine (HYP) and lactate (LAC) by the PFKP rs2388595 SNP and other factors (**A**). Proteomics of CD235^+^ CD71, WBC-/PLT-/reticulocyte-depleted, mature RBC-enriched samples (schematic in **B**; flow-diagrams in **C**) confirmed that PFKP accounts for ∼10% of the RBC PFK isoforms (**D**). In **E**, total abundance (by proteomics-derived integrated peak area, arbitrary units) for glucose/monocarboxylate transporters and glycolytic enzymes (official gene symbols) in RBCs as a function of PFKP rs2388595 alleles (n=643; storage day 42). In **F**, weekly measurements of glycolytic metabolites in RBCs (from day 0 to 42; n=6). In **C**, storage experiments in presence of 1,2-^13^C_2_-glucose (^13^C-labeled isotopologues as shown as % of the total levels of each metabolite – **G**).

Since PFKP is a platelet (PLT)-specific isoform, we questioned whether our proteomics measurements and mQTL results had been influenced by changes in cellular blood counts and PLT contamination. However, we found no association between ATP measurements and RBC or PLT counts at the time of donation, nor between PFKP SNP or PFKP protein levels and PLT counts (**Supplementary Figure 7.A-C**). We then isolated ultra-pure mature RBC populations from 5 independent volunteers, whose blood was leukofiltered (log2.5 and log4 PLT and WBC depletion, respectively), and flow-sorted to remove residual WBCs, PLTs and immature reticulocytes (based on CD41, CD45, CD62P, and CD71 cell surface markers), before positive flow-sorting-based enrichment for CD235a^+^ CD71^−^ mature RBCs (**Figure 6.B-C**). Proteomics analyses of the ultra-pure mature RBC samples still identified PFKP, which comprised ∼11% of the total PFK isozyme population – ∼89% of which was represented by PFKL/M (**Figure 6.D**). This analysis also confirmed the presence of CD38 in ultra-pure mature RBCs. The RBC proteome did not change significantly in the stored RBC samples from the recalled donors, neither as a function of storage or PFKP allele dosage (**Supplementary Figure 7.D**).

### Understanding the role of PFKP in stored RBCs through dynamic modeling

RBCs from donors carrying two alleles of the rs2388595 PFKP SNP had ∼30% higher levels of PFKP (**Figure 6.E**), significantly higher levels of GAPDH, and higher medians (albeit not significant) for LDHB (**Figure 6.E**). To understand changes in glycolysis as a function of storage duration, we combined time course measurements at day 10, 23 and 42 from recalled donor units (n=643 – **Supplementary Figure 6.F**), and data from a separate storage experiment using 6 new pRBCs units; in the latter study, units were sterilely sampled weekly to obtain a more granular understanding of the progression of the metabolic storage lesion to glycolysis (**Figure 6.F**). Altogether, these results indicated a drop in ATP, down to 20% of the initial levels by storage day 42. Storage was accompanied by progressive declines in total NAD pools (threefold decrease), while nicotinamide levels increased 2.55 fold (**Figure 6.F**), similar to prior reports.^26,40^ In a separate experiment, pRBC units were spiked with 1,2-^13^C_2_-glucose (50% of total), to trace ^13^C incorporation into glycolytic metabolites (**Figure 6.G**). These combined analyses show that: (i) glycolytic fluxes are active during storage; (ii) when 50% of glucose is labeled, ∼40% of lactate is labeled by the end of storage (estimates adjusted based on the consideration that only 50% of 1,2-^13^C_2_-glucose-derived lactate can be labeled), consistent with the literature^19,42^; (iii) de novo BPG synthesis is insufficient to maintain the BPG pools; and finally, (iv) the PFK-catalyzed step appears to be rate-limiting, since FBP declines at the fastest rate of all glycolytic metabolites despite ongoing synthesis (**Figure 6.F-G**).

To understand the role of the PFKP isozyme in human RBCs, we constructed a mass action stoichiometric simulation model of the in vivo RBC glycolytic pathway at steady state.^49,50^ Differences in kinetic and regulatory properties suggest distinct metabolic roles for PFK isozymes, ^51,52^ whose simultaneous expression favors metabolic plasticity in response to various conditions. In particular, PFKP is notable among PFK isozymes as it is not allosterically activated by either fructose 1,6-bisphosphate or glucose 1,6-bisphosphate. Furthermore, PFKP has the lowest affinity for ATP as a substrate and a relative insensitivity to inhibition by ATP, suggesting that it would have a physiologically relevant role when ATP concentration is low.

First, we replaced mass action rate laws with detailed enzyme kinetic equations and parameters for the key regulatory enzymes hexokinase, phosphofructokinase, and pyruvate kinase^23,53^, ensuring their regulatory properties were represented in the model.

Modeling isozymes is generally associated with difficulties trying to ascertain the relative contributions of each isozyme within a cellular context^54^. Difficulties in discerning differences in catalytic and regulatory properties are further complicated when quaternary structure can be comprised from heterologous subunit oligomerization, as illustrated by PFK. Thus, we built two glycolytic pathway models, one containing only comprised of PFK liver/muscle (PFKL/M) isozymes, and one containing both PFKL/M and PFKP, to gain qualitative insight into the underlying mechanisms responsible for the association of ATP levels with PFKP. Glycolytic enzyme abundances were estimated based on proteomics data for donors carrying 0 or 2 alleles of PFKP SNP rs2388595 (**Supplementary Figure 7.E**). We then assumed that the existing PFK reaction was catalyzed by the tetrameric enzyme formed by liver and muscle isozymes in erythrocytes – the most abundant isoforms based on our proteomics analyses (∼89%). In the first model, PFK abundance was set to 1e-7 M^55^; in the second model, PFKP was set to account for ∼11% of the total PFK isozyme. Mature RBCs do not contain significant concentrations of fructose 2,6-bisphosphate^19,22,27,42^; therefore, we assumed that fructose 2,6-bisphosphate is negligible as an effector in the intracellular RBC environment. Finally, we searched the literature and parameterized the model based on published enzyme kinetics and parameters.^51,56–58^. We then simulated and updated models, checking the eigenvalues of the Jacobian matrix to ensure a stable homeostatic state was reached (**Supplementary Material**).

Upon transfusion, stored RBCs must recover their ATP content to restore their biophysical properties (e.g., stiffness^59^), as the “energy-less RBC is rapidly lost”^8^ via extravascular hemolysis^60^. We reasoned that PFKP may play a role in the initial post-transfusion recovery of cellular ATP.

Therefore, models representing RBC glycolysis with and without PFKP were simulated after a sudden 30% decrease in ATP concentration with a concomitant ADP increase– similar to changes observed in RBCs stored up to the seventh day, before significant ATP breakdown to hypoxanthine occurs.^26^ Glycolytic fluxes and metabolite concentrations were then modeled over a time scale of 10^−4^ to 10^4^ hours prior to and after the simulated drop in ATP (**Supplementary Figure 7.F-H**). Models representing RBC glycolysis in donors expressing higher levels of PFKP (i.e., donors carrying 2 alleles of the rs2388595 SNP) were simulated in an in vivo-like environment (**Supplementary Figure 7.F**). Simulations revealed that dynamic response of glycolysis was strongly influenced by changes in the adenylate phosphates (AMP, ADP, ATP – AXP). Phase planes of PFKL/M and PFKP fluxes vs adenylate phosphates demonstrate decreased PFKP and increased PFKL/M deviation from the physiological steady state in donors expressing higher levels of PFKP (i.e., donors carrying 2 alleles of the rs2388595 SNP), suggesting an increased capacity maintain glycolytic flux.

The sudden drop of ATP reduced its regulatory influence as an allosteric inhibitor, and the subsequent influx of AMP from the salvage pathway activated the PFK enzymes on the fast time scale. The simulation shows that the adenylate kinase initially redistributes the phosphates as AMP enters the system, and ATP is formed by lower glycolysis. Though PFKL/M and PFKP are both activated by AMP, PFKL/M will experience more inhibitory effects due to ADP as compared to PFKP. However, as F6P depleted, F1,6BP is replenished, and ATP is recovered, PFKL/M returned back to their steady states (**Supplementary Figure 7.G-H**). As the flux of the PFK isoforms return to steady state levels, the excess AMP in the system flowed out of the system.

The existence of PFKP boosted the initial recovery response and prevented the overshoot of PFKL/M when the model returned to its physiological steady state, a likely result of differences in regulation as PFKP switched between low-flux and higher-flux states. Although simplifying assumptions for parameter reconciliation necessitates caution in numerical interpretation of the simulation, the observed dynamic responses are consistent with previous *in silico* and experimental kinetic studies in other cellular systems,^51,52^ suggesting that kinetic and regulatory properties of PFKP play a role in preventing loss of the total AXP pool.

### ATP and hypoxanthine are biomarkers of hemolysis in vitro and in vivo

End-of-storage spontaneous hemolysis ex vivo was measured in 12,753 units from the REDS RBC Omics index donor cohort (**Figure 7.A**). High levels of ATP and glycolytic metabolites in cluster 4 were associated with significantly lower levels of hemolysis (**Figure 7.A**), as were units from donors carrying two alleles for the rs2388595 SNP (**Figure 7.B**). However, spontaneous hemolysis was a poorly sensitive marker of unit quality, since >97% of units (independent of cluster or genotype) had <1% hemolysis. As such, we focused on the other quality gold standard, PTR. Specifically, we determined metabolic correlates to PTR in a separate cohort of 79 volunteers from the Donor Iron Deficiency Study (DIDS) of frequent donors (**Figure 7.C**).^61^ From this analysis, glycolytic metabolites and ATP and hypoxanthine levels were positively and negatively correlated with PTR, respectively, in healthy volunteers receiving autologous ^51^Cr-labeled RBCs (**Figure 7.C**). By accessing the “vein-to-vein” database of the REDS RBC Omics program, we first linked blood donor PFKP SNPs to decreases in adjusted hemoglobin increments as a function of allele frequency for rs2388595 (reverse linkage in the discordant rs61835134 SNP – **Data S1**), in patients receiving single irradiated RBC units (**Figure 7.D**). We then linked ATP and hypoxanthine levels to hemoglobin increments (n=5,816 records available in the database) and bilirubin increments (n=914) upon transfusion of heterologous products from REDS donors to ill recipients requiring transfusion (**Figure 7.E**). Results indicate a significant positive association between ATP levels and hemoglobin increments in transfusion recipients, along with decreased bilirubinemia (i.e. lower hemolysis in vivo) in patients receiving these units. In addition, the opposite results were observed as a function of hypoxanthine levels, yielding lower hemoglobin increments and higher bilirubin increments (i.e., higher in vivo hemolysis – **Figure 7.E-F**).

**Figure 7.**
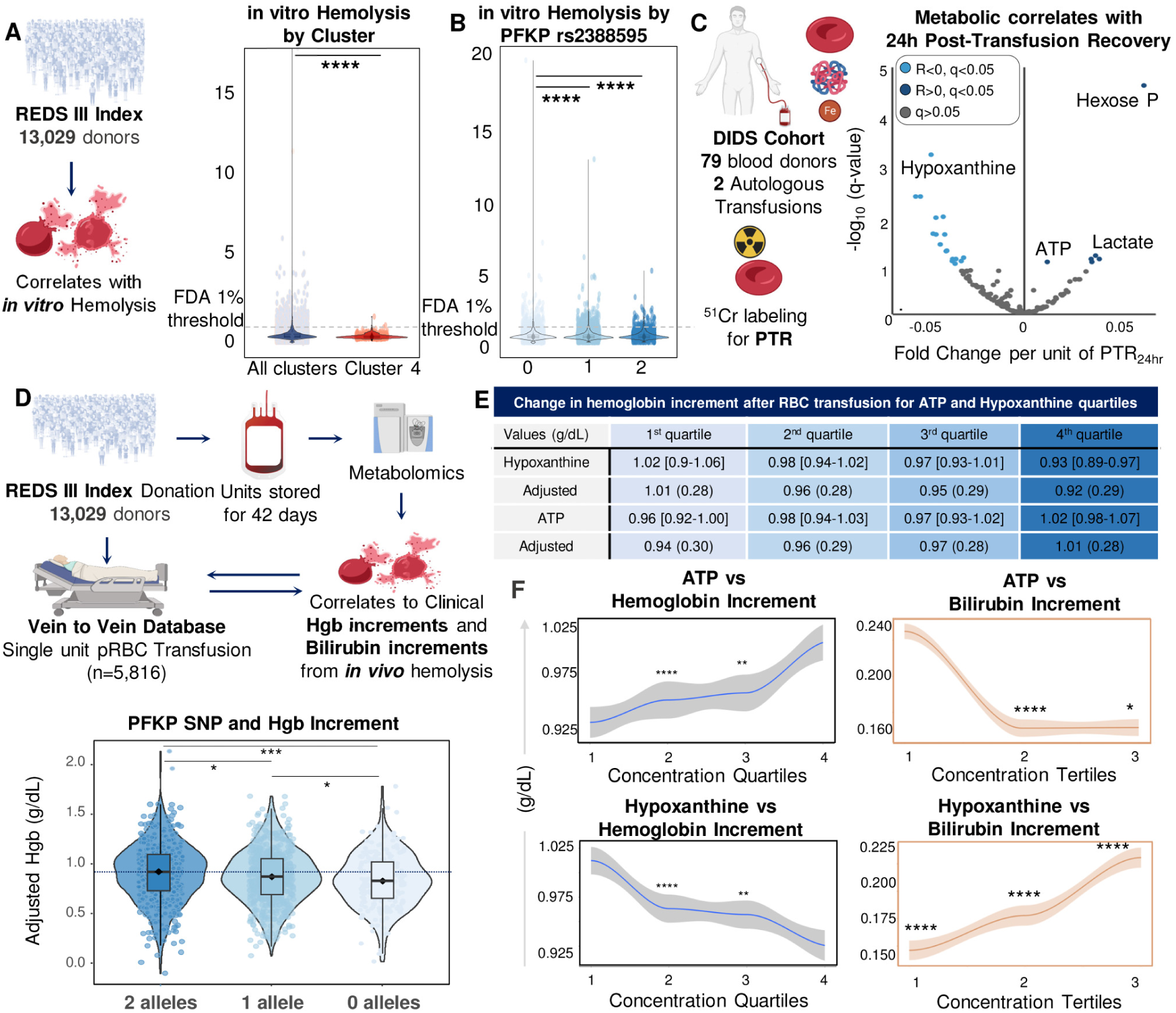
– PFKP SNP, ATP and hypoxanthine are markers of in vitro and in vivo hemolysis in healthy and critically ill individuals receiving autologous and non-autologous transfusion, respectively. Compared to the rest of the population, units from donors in cluster 4 (**A**) or donors carrying two alleles of the PFKP SNP 2388595 (**B**) showed significantly lower levels of end of storage in bag hemolysis. In a separate cohort of 79 healthy volunteer donations and autologous transfusion events, glycolytic metabolites and ATP and hypoxanthine levels were positively and negatively associated to post-transfusion recovery with ^51^Cr in healthy autologous volunteers, respectively (**C**). Association between adjusted hemoglobin increments in clinical recipients of irradiated single pRBC unit transfusions and blood donor PFKP status for rs2388595 (0 alleles n=521; 1 allele, n=576; 2 alleles, n=204) (**D**). End of storage ATP and hypoxanthine levels are associated to hemoglobin (n=5,816 records available in the “vein-to-vein” database) and bilirubin increments (n=914) upon heterologous transfusion of products from REDS donors to critically ill recipients requiring transfusion (**E-F**).

## DISCUSSION

Herein we detail how energy metabolism of stored human and murine RBCs is impacted by biological (age, sex), processing (additives) and genetic factors, and in turn influences RBC hemolysis in vitro and in vivo. Specifically, we identified an association between elevated glycolysis and common polymorphisms in PFK (PFKP more significantly than PFKM) and HK1, two well-established rate-limiting enzymes of glycolysis.^15^ While PKFP SNPs were previously associated with PLT counts in the GWAS catalogue,^47^ proteomics analyses of ultra-pure RBC populations here showed that PFKP is expressed in mature RBCs, accounting for ∼11% of total PFK isoforms. Lactate levels were also associated with PFKM SNPs, such as the missense mutation Gln76Lys (rs4760682), although with a much lower significance, likely owing to the lower minor allele frequency of this variant in the REDS donor population.

RBC units from these donors with elevated end-of-storage ATP levels were primarily stored in AS-3, an additive containing high levels of citrate to compensate for the lack of mannitol as an osmoprotectant. This formulation was designed to mitigate osmotic stress that lead to elevations of intracranial pressure in recipients, making this additive appropriate for pediatric patients and patients with traumatic (brain) injury.^40^ The excess citrate in AS-3 could play a hitherto unappreciated role in the economy of RBC glycolytic fluxes. Previous studies showed that RBCs possess functional cytosolic isoforms of Krebs cycle enzymes that could utilize citrate as a substrate to contribute to NADPH homeostasis (e.g., isocitrate dehydrogenase 1); enzymes in this pathway could also contribute to oxaloacetate and malate pools (via cytosolic malate dehydrogenase 1 and acetyl-coA lyase),^19,27,62,63^ ultimately feeding back into late glycolysis at the PEP step by a PEP carboxylase-like mechanism mediated by hemoglobin.^64^ PFK inhibition by citrate is a well-established concept in cells endowed with mitochondria, a process that inhibits glycolysis as a function of the abundance of fatty acid oxidation–derived citrate – the Randle cycle.^65^ However, there are marked dose-response isoform-specific effects for PFKM (IC_50_^citrate^ 982L±L109 µM) and PFKP (IC_50_^citrate^ 177L±L132 µM), but not PFKL.^51^ Since normal plasma citrate concentrations in humans are within a range of ∼100–150 µM^66^, citrate-dependent isoform-specific regulatory effects on PFK may not only be relevant to the supra-physiological conditions of anticoagulant-supplemented storage additives (up to 25 mM citrate in AS-3), but also in vivo. This may also have implications for considering citrate as a regulator of immune^67^ and cancer^68^ cell metabolism, at least in subjects carrying one or two alleles of the PFKP SNPs, i.e., 15% of the donor population in the present study.

In this view, it is interesting to note that, when specific activities are considered, it has been reported that citrate may no longer act as an inhibitor, but rather as activator of PFK when ATP levels are low (sub-physiological), such as those observed in end-of-storage RBCs.^51^ Such an activating effect is also isoform specific, with PFK-PL>LPFK-LL>LPFK-M.^51^ Our data and the simulation of glycolysis fit with this model, whereby SNPs (e.g., rs2388595) associated with higher PFKP expression levels are also associated with higher glycolytic fluxes in stored human or murine RBCs, but not in fresh murine RBCs. In-depth quantitative modeling of glycolysis, including modeling of how glycolysis responds to perturbation of the ATP/ADP ratio with a fixed total AXP pool, did not reveal an obvious distinguishing metabolic feature of cells with high or low PFKP that would be explained by the observed PFKP genetics. One possible interpretation is that PFKP-dependent effects relate to preventing loss of the total AXP pool, although further work is needed to test this hypothesis.

Translationally, as the field is evaluating the implementation of Precision Transfusion Medicine arrays^69^ in routine blood donor screening (e.g., to identify blood group genotypes and minor/rare antigens), monitoring the rs2388595 or rs61835134 PFKP SNPs could identify donors whose blood would store and perform better after transfusion. Since big leaps in ex vivo generation of blood products have been documented in recent years,^70^ the identification of genetic polymorphisms that could maximize the quality and post-transfusion performances of ex vivo generated RBCs holds important translational implications.

We also identified an association between polymorphisms in CD38 and BST1 – both of which are ADP-ribosyl cyclases (1 and 2) – and NAD(P), PPP, and late glycolysis in both fresh and stored human and murine RBCs. Our data suggests a role for CD38/BST1 in NAD(P) catabolism in stored RBC units as a fuel for ATP synthesis, upon phosphorolysis of ADP-ribose and re-entering of pentose phosphate into glycolysis downstream of glyceraldehyde 3-phosphate.

In conclusion, our findings provide “real-world” clinical data on the potential utility of ATP and hypoxanthine levels – and the genetic traits linked to them – as markers of stored RBC hemolytic propensity, both in the bag and, most importantly, in vivo following transfusion.^71^ To this end, we combined autologous PTR studies on an independent cohort with clinical data by interrogating a “vein-to-vein” database containing records for thousands of transfusion events of units from REDS RBC Omics donors.^72^ Accumulation of hypoxanthine also informs on both the energy and redox status of stored RBCs, as ATP consumption and deamination are favored by BPG consumption and oxidant stress, factors that contribute to the enzymatic activity of AMPD3^24,25^. Current gold standards of RBC storage quality, in bag hemolysis and PTR, are not routinely tested for all issued blood products, rather they are only measured when securing FDA or European Council approval for newly licensed blood products. Here we show that stored RBC ATP and hypoxanthine levels are not only relevant surrogates for these measures, but are also amenable to ultra-high-throughput screening at scale, and could potentially be implemented for all of the >100 million RBC units donated worldwide. Since RBC ATP levels are also essential regulators of oxygen kinetics,^73^ our results represent a step forward towards identifying markers that can not only inform on the quality of the stored products but also, potentially, predict their efficacy following transfusion.

## Limitations of the study

This study focuses on RBC metabolism, arguably the simplest eukaryotic cell; therefore, these results may not necessarily extend to more complex cell types. While glycolysis is the sole energy generating pathway in mature RBCs, other pathways like redox homeostasis via glutathione metabolism may be equally relevant in the economy of RBC storage quality and/or post-transfusion performances. One major limitation of mQTL studies is that genetically conserved, non-polymorphic regions would not be highlighted. In addition, since our studies are based on volunteers who are sufficiently healthy to donate blood, genetic polymorphisms that are either incompatible with life or associated with major hematological effects would not be captured by our analysis. As such, signals from rare polymorphisms would not be detected unless additionally diverse populations (e.g., non-healthy, non-blood donors, other ancestries) were also investigated. Nonetheless, we did detect signals for genes like G6PD, PKLR or ADA, all of which are associated with hematological conditions, suggesting that signatures for these genotypes are still detectable in the metabolome of RBCs from otherwise asymptomatic heterozygous carriers who donate blood, prompting considerations on the very definition of “health” at population scale. Despite strong effects of additives, sex, age and genetic ancestry, a large percentage of ATP variance remains unexplained, with no significant interactions observed between genotypes and biological factors, suggesting that other factors (e.g., exposome) may contribute to RBC glycolysis. Yet, genetic polymorphisms and ATP levels were linked to relevant in vitro and in vivo hemolysis post-transfusion outcomes. Finally, *in silico* modeling suggests that PFKP-dependent effects may relate to preventing loss of the total AXP pool, although more refined models are necessary to elucidate precise mechanisms.

## Acknowledgements

This study was supported by funds by the National Heart, Lung, and Blood Institute R21HL150032, R01HL146442, R01HL149714 (to AD and JCZ), R01HL148151 (to AD, SLS and JCZ) and R01HL126130 (NR). The authors acknowledge support from the REDS RBC Omics and REDS-IV-P CTLS programs, sponsored by the National Heart, Lung, and Blood Institute contract 75N2019D00033, and from the NHLBI Recipient Epidemiology and Donor Evaluation Study-III (REDS-III) RBC Omics project, which was supported by NHLBI contracts HHSN2682011-00001I, –00002I, –00003I, –00004I, –00005I, –00006I, –00007I, –00008I, and – 00009I. The autologous post-transfusion recovery studies were funded by NHLBI R01HL133049. The content is solely the responsibility of the authors and does not necessarily represent the official views of the National Institutes of Health.

## Authors’ contributions

TN, DS, AD performed metabolomics analyses. TN, DS, JAR, AD performed method development, quality control and metabolomics data analysis. EE, AM, GPP performed human mQTL analyses; GRK, GAC performed mQTL analyses for the Diversity Outbred mice; AH, JCZ performed the Diversity Outbred mouse studies; MD, KEH, KCH performed proteomics studies; ZH and BOP generated the in silico simulation of glycolysis; SLS, EAH performed post-transfusion recovery studies (DIDS cohort); CE, AK, XD, EE, GRK, GAC, NR, AD performed statistical analyses; NR performed linkage analyses to the vein-to-vein database; MS, PJN, SK, MPB performed and supervised the REDS RBC Omics study; TN, EE, GRK, AK, CE, AD prepared figure panels; AD wrote the first version of the manuscript, that was critically reviewed and approved by all authors.

## STAR METHODS

### RESOURCE AVAILABILITY

#### Lead contact

Further information and requests for resources and reagents should be directed to and will be fulfilled by the Lead Contact, Angelo D’Alessandro

#### Materials availability

No new materials were generated as part of this study. Information on the human and murine samples tested in this study is detailed below.

#### Data and code availability

- Human GWAS summary statistics are available through the GWAS catalogue with the following identifiers: 2,3-Phosphoglycerate – GCST90421022; 2,3-Bisphosphoglycerate – GCST90421023; ATP – GCST90421024; Fructose bisphosphate – GCST90421025; Glucose – GCST90421026; Glyceraldehyde 3-phosphate – GCST90421027; Hexose phosphate – GCST90421028; Lactate – GCST90421029; NADPplus – GCST90421030; Phosphoenolpyruvate – GCST90421031; Pyruvate – GCST90421032. Processed J:DO mouse data and results are uploaded to the QTLViewer webtool (https://churchilllab.jax.org/qtlviewer/Zimring/RBC;^74^) for interactive analyses and download. Omics data sets and their elaborations are available in the supplementary tables.
- All data and code for the analysis pipeline are available at figshare (https://doi.org/10.6084/m9.figshare.24456619) and data S1.
- Any additional information required to reanalyze the data reported in this paper is available from the Lead Contact upon request.

## EXPERIMENTAL MODEL AND SUBJECT DETAILS

### Donor recruitment in the REDS RBC Omics study

#### Index donors

A total of 13,758 donors were enrolled in the Recipient Epidemiology and Donor evaluation Study (REDS) RBC Omics at four different blood centers across the United States (https://biolincc.nhlbi.nih.gov/studies/reds_iii/) after obtaining informed consent. Demographic information is provided in Data S1. Of the donors, 97% (13,403) provided informed consent and 13,029 were available for metabolomics analyses in this study – henceforth referred to as “index donors”. A subset of these donors (n=12,753) was evaluable for hemolysis parameters, including spontaneous hemolysis in ∼42-day stored leukocyte-filtered packed RBCs derived from whole blood donations from this cohort ^38^.

#### Recalled donors

A total of 643 donors scoring in the 5^th^ and 95^th^ percentile for hemolysis parameters at the index phase of the study were invited to donate a second unit of pRBCs, a cohort henceforth referred to as “recalled donors”. These units were assayed at storage days 10, 23 and 42 for hemolytic parameters and mass spectrometry-based high-throughput metabolomics ^75^, proteomics ^76^, lipidomics ^77^ and ICP-MS analyses ^78^. Under the aegis of the REDS-IV-P project ^79^, a total of 1,929 samples (n=643, storage day 10, 23 and 42) were processed with this multi-omics workflow. Extracellular vesicles were measured by flow-cytometry^80^.

### Vein-to-vein database: General Study Design

We conducted a retrospective cohort study using electronic health records from the National Heart Lung and Blood Institute (NHLBI) Recipient Epidemiology and Donor Evaluation Study-III (REDS-III) program available as public use data through BioLINCC ^81,82^. The database includes blood donor, component manufacturing, and patient data collected at 12 academic and community hospitals from four geographically diverse regions in the United States (Connecticut, Pennsylvania, Wisconsin, and California) for the 4-year period from January 1, 2013 to December 31, 2016. Genotype and metabolomic data from the subset of blood donors who participated in the REDS-III RBC-Omics study ^83^ was linked to the dataset using unique donor identifiers. Data regarding blood donor demographics (e.g., age, sex, body mass index, ABO/Rh), collection date and methods (e.g., whole blood or apheresis), as well as component manufacturing characteristics (e.g., timing of leukoreduction, additive solution, irradiation status) were extracted for each RBC unit collected. Available donor genetic polymorphism and metabolomic data was linked to issued RBC units using random donor identification numbers. Among transfusion recipients, we included all adult patients who received a single RBC unit during one or more transfusion episodes between January 1, 2013 and December 30, 2016. Recipient details included age, sex, body mass index (BMI), along with storage age, and blood product issue date and time. We collected hemoglobin and bilirubin levels measured by the corresponding clinical laboratory prior to and following each RBC transfusion event.

### Donor recruitment in the Donor Iron Deficiency Study (DIDS)

79 repeat blood donors were enrolled as part of the DIDS study for baseline and post-iron repletion before determination of autologous post-transfusion recovery (PTR) studies of day 42 stored units with ^51^Cr, as described ^61^.

### Diversity outbred mice

Mouse storage studies utilized a population of 525 diversity uutbred mice obtained from the Jackson Laboratory (J:DO) derived from extensive outbreeding of eight inbred founder strains that represent genetically distinct lineages of the house mouse ^84,85^. All animal procedures were approved by the University of Virginia IACUC (protocol no. 4269).

## METHOD DETAILS

Methods for the determination of FDA-standard spontaneous (storage) hemolysis test, osmotic hemolysis (pink test) and oxidative hemolysis upon challenge with AAPH have been extensively described^86^.

### Storage hemolysis

After storage under routine blood bank conditions (1-6 °C) for 39 to 42 days, the contents of each unit was transferred into a 15 mL conical tube from which 2 aliquots (1 mL each) were processed for the hemolytic assays. One aliquot was used for the quantification of spontaneous storage hemolysis and the other for the stress-induced hemolysis assays. Percent end-of-storage hemolysis was determined according to the following equation:

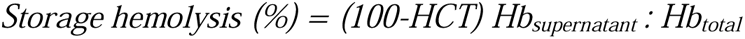

Sample hematocrit (HCT) was determined by collecting blood samples into capillary tubes, which were centrifuged in a micro-HCT centrifuge (LW Scientific, Lawrenceville, GA). Hb_supernatant_ refers to the levels of free hemoglobin obtained after centrifugation (1500 g, 10 min, 18 °C) measured in the supernatant. Hb_total_ refers to the total amount of sample hemoglobin before centrifugation. In the entire study, hemoglobin concentrations (micromolar) were determined by Drabkin’s method.^87^

For the evaluation of stress-induced hemolysis, stored RBCs were washed (1500 g, 10 min, 18 °C) 3 times with phosphate-buffered saline (PBS) to remove plasma and additive solution, and immediately subjected to osmotic or oxidative stress assays.

### RBC osmotic hemolysis

Osmotic hemolysis was determined by a modified pink test assay previously used for diagnosis of genetic mutations that affect RBC osmotic fragility (eg, sickle cell disease, thalassemia, and spherocytosis) ^88,89^. Washed RBCs were incubated under static conditions (4 hours at 22 °C) in pink test buffer (a hypotonic Bis-Tris buffer containing 25 mM sodium chloride, 70 mM 2,2-bis(hydroxymethyl)-2,2′,2″nitrilotriethanol (Bis-Tris) buffer, and 135 mM glycerol; pH 6.6) at a final concentration of 1.6% ± 0.2% after which samples were centrifuged (1500 g, 10 min, 18 °C), and percent osmotic hemolysis was determined:

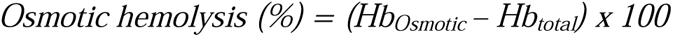

for which Hb_osmotic_ corresponds to supernatant cell-free hemoglobin of pink test–treated RBCs, and Hb_total_ refers to the total amount of hemoglobin in each sample.

### RBC oxidative hemolysis

Testing for donor differences in RBC susceptibility to oxidative hemolysis was performed as described by Kanias et al.,^86^ by incubating RBCs in the presence of 2,2′-azobis-2-methyl-propanimidamide dihydrochloride (AAPH; 150 mM). Thermal (37 °C) decomposition of AAPH generates peroxyl radicals leading to lipid peroxidation–mediated hemolysis. Washed RBCs were suspended with PBS to a final concentration of 3.5% ± 0.5%. Aliquots (0.21 mL) were transferred into microplates to which 0.09 mL of AAPH (0.5 M) or PBS was added. The plates were incubated (37 °C) under static conditions for 1.5 hours, after which the plates were centrifuged (1500 g, 10 min, 18 °C), and AAPH-induced oxidative hemolysis was determined by using the following formula:

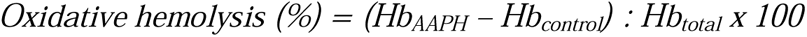

where Hb_AAPH_ corresponds to supernatant cell-free hemoglobin of AAPH-treated RBCs, Hb_control_ corresponds to supernatant cell-free hemoglobin from untreated RBCs, and Hb_total_ refers to the total amount of hemoglobin in each sample.

### Storage with weekly sampling in presence or absence of 1,2-^13^C_2_-glucose

Whole blood units were donated by 6 healthy donor volunteers in CP2D (Haemonetics, Boston, MA, United States) and suspended in AS-3 additive solution after leukofiltration and plasma removal. Sterile weekly sampling was performed from day 0 to day 42 prior to metabolomics analyses. In a separate storage experiment, six whole blood units were processed as above and suspended in a custom AS-3 formulation containing 50%L1,2-^13^C_2_-Glucose (Sigma Aldrich, St. Louis, MO, United States). When calculating % ^13^C enrichment as a fraction of the total isotopologues. Please, note that only 50% of triose phosphate and downstream glycolytic metabolites would contain 1,2-^13^C-glucose-derived heavy C, and as such % isotopologue estimates for triose compounds were adjusted accordingly.

### High-throughput metabolomics

Metabolomics extraction and analyses in 96 well-plate format were performed as described.^90,91^ RBC samples were transferred on ice to 96 well plates and frozen at –80 °C at Vitalant San Francisco prior to shipment on dry ice to the University of Colorado Anschutz Medical Campus. Plates were thawed on ice then a 10 µL aliquot of each sample was transferred with a multi-channel pipettor to 96-well extraction plates. A volume of 90 µL of ice cold 5:3:2 MeOH:MeCN:water (*v/v/v*) was added to each well with an electronically-assisted cycle of sample mixing repeated three times. Extracts were transferred to 0.2 µm filter plates (Biotage) and insoluble material was removed under positive pressure using nitrogen applied via a 96-well plate manifold. Filtered extracts were transferred to an ultra-high-pressure liquid chromatograph (UHPLC, Vanquish) equipped with a plate charger. A blank containing a mix of standards detailed before^92^ and a quality control sample (the same across all plates) were injected 2 and 4 times per plate, respectively, to qualify instrument performance throughout the analysis. For each measured metabolite, median values on each plate were determined; all samples were normalized intra-plate to the respective median value prior to merging the entire dataset (principal component analyses – PCA of pre-vs post-normalization data are shown in **Data S1**). Metabolites were resolved on a Phenomenex Kinetex C18 column (2.1 x 30 mm, 1.7 µm) at 45 °C using a 1-minute ballistic gradient method in positive and negative ion modes (separate runs) exactly as previously described^90^. The UHPLC was coupled online to a Q Exactive mass spectrometer (Thermo Fisher). The Q Exactive MS was operated in Full MS mode (2 μscans) from 60 to 900 m/z at 70,000 resolution with 4 kV spray voltage, 45 sheath gas, 15 auxiliary gas. Following data acquisition, .raw files were converted to .mzXML using RawConverter then metabolites assigned and peaks integrated using ElMaven (Elucidata) in conjunction with an in-house standard library ^93^. A list of experimental m/z, retention times, metabolite IDs (either KEGG or HMDB) and adducts is provided in **Data S1**.

### High-throughput proteomics

Proteomics analyses were performed as described.^43,94^ A volume of 10 μL of RBCs were lysed in 90 μL of distilled water; 5 μL of lysed RBCs were mixed with 45 μL of 5% SDS and then vortexed. Samples were reduced with 10 mM DTT at 55 °C for 30 min, cooled to room temperature, and then alkylated with 25 mM iodoacetamide in the dark for 30 min. Next, a final concentration of 1.2% phosphoric acid and then six volumes of binding buffer (90% methanol; 100 mM triethylammonium bicarbonate, TEAB; pH 7.1) were added to each sample. After gentle mixing, the protein solution was loaded to a S-Trap 96-well plate, spun at 1500 x g for 2 min, and the flow-through collected and reloaded onto the 96-well plate. This step was repeated three times, and then the 96-well plate was washed with 200 μL of binding buffer 3 times. Finally, 1 μg of sequencing-grade trypsin (Promega) and 125 μL of digestion buffer (50 mM TEAB) were added onto the filter and digested carried out at 37 °C for 6 h. To elute peptides, three stepwise buffers were applied, with 100 μL of each with one more repeat, including 50 mM TEAB, 0.2% formic acid (FA), and 50% acetonitrile and 0.2% FA. The peptide solutions were pooled, lyophilized, and resuspended in 500 μL of 0.1 % FA. Each sample was loaded onto individual Evotips for desalting and then washed with 200 μL 0.1% FA followed by the addition of 100 μL storage solvent (0.1% FA) to keep the Evotips wet until analysis. The Evosep One system (Evosep, Odense, Denmark) was used to separate peptides on a Pepsep column, (150 µm inter diameter, 15 cm) packed with ReproSil C18 1.9 µm, 120A resin. The system was coupled to a timsTOF Pro mass spectrometer (Bruker Daltonics, Bremen, Germany) via a nano-electrospray ion source (Captive Spray, Bruker Daltonics). The mass spectrometer was operated in PASEF mode. The ramp time was set to 100Lms and 10 PASEF MS/MS scans per topN acquisition cycle were acquired. MS and MS/MS spectra were recorded from *m/z* 100 to 1700. The ion mobility was scanned from 0.7 to 1.50LVs/cm^2^. Precursors for data-dependent acquisition were isolated withinL±L1LTh and fragmented with an ion mobility-dependent collision energy, which was linearly increased from 20 to 59LeV in positive mode. Low-abundance precursor ions with an intensity above a threshold of 500 counts but below a target value of 20000 counts were repeatedly scheduled and otherwise dynamically excluded for 0.4Lmin.

### Preparation of pure mature RBCs

Whole blood was obtained from 6 healthy donor volunteers at Columbia University in New York, NY, USA, upon signing of informed consent. Whole blood from 5 healthy volunteers was collected into EDTA tubes. Blood was spun at 800 g for 10 minutes, plasma was removed, and 50 µL of packed RBCs from each donor was combined prior to leukoreduction via filtration (Pall Medical) to remove log4 (99.99%) and log2.5 residual white blood cells (WBCs) and platelets (PLTs), respectively, as standard workflow in modern blood banks in most US states.^95^. The mixed RBCs were washed in PBS and 200 µL of packed RBCs was resuspended in 10mL PBS and stained with antibodies CD45 PE-Cy7 (1:400), CD41 APC-Cy7 (1:400), CD235a BV605 (1:1000), and CD71 PE (1:100) for 20 minutes at room temperature. Samples were washed with PBS and resuspended to a final volume of 1 mL. Immediately before sorting, 30 µL of RBCs was diluted into 10mL PBS supplemented with 20 µL thiazole orange. Purified RBC populations were achieved using BD FACSAria cell sorters. CD45+ white blood cell and CD41+ platelets were excluded from RBC singlets and reticulocytes (CD235a+TO+) and mature RBCs (CD23a+CD71-TO-) were sorted. Aliquots of sorted cells were re-analyzed on the cell sorter to determine purity, indicating >99.99% mature RBCs.

### mQTL analysis in REDS RBC Omics

The workflow for the mQTL analysis of ATP, hypoxanthine and glycolytic metabolites is consistent with previously described methods from our pilot mQTL study on 250 recalled donors.^96^ Details of the genotyping and imputation of the RBC Omics study participants have been previously described by Page, et al. ^29^ and Nemkov et al. ^43^ Metabolites with missing data and zeros were both treated identically. We separated the participant data by blood storage additive type and excluded metabolites with greater than 10% missingness from each additive set, respectively. Relatively quantified metabolites were natural log transformed. These groups of metabolites then had missing metabolite levels imputed using QRILC^97^, implemented in the R package QRILC^97^. After imputation, all metabolites were inverse normal transformed using the R package GenABEL rntransform command^98^. Metabolites with more than 10% missing were not considered in these analyses. Briefly, genotyping was performed using a Transfusion Medicine microarray ^69^ consisting of 879,000 single nucleotide polymorphisms (SNPs); the data are available in dbGAP accession number phs001955.v1.p1. Imputation was performed using 811,782 SNPs that passed quality control. After phasing using Shape-IT ^99^, imputation was performed using Impute2 ^100^ with the 1000 Genomes Project phase 3 ^100^ all-ancestry reference haplotypes. We used the R package SNPRelate ^101^ to calculate principal components (PCs) of ancestry. We performed association analyses for glycolytic metabolites using an additive SNP model in the R package saigeGDS^102^ and 13,091 study participants who had both metabolomics data and imputation data on serial samples from stored RBC components that passed respective quality control procedures. The full cohort was stratified into four groups based on inferred genetic ancestry: African American (AFR, N=1,503), East and South Asian (ASN, N=1,560), European (N=7,943), and admixed Hispanic (HIS, N=986). Multivariate regressions were performed within each of these groups, adjusting for sex, age (continuous), frequency of blood donation in the last two years (continuous), blood donor center, and ten genetic PCs. Results from each group were meta-analyzed using a mixed random effect regression using the tool *random-METAL* (https://github.com/explodecomputer/random-metal). Statistical significance was determined using a p-value threshold of 5×10^−8^. We only considered variants with a minimum minor allele frequency of 1% and a minimum imputation quality score of 0.80. We only considered variants with a minimum minor allele frequency of 1% and a minimum imputation quality score of 0.80. Associated SNPs were considered independent when exhibiting linkage of r^2^ < 0.6. Conditional regressions were performed to identify additional independent genome-wide associated SNPs for each metabolite. The ENSEMBL Variant Effect Predictor^103^ was used to annotate the top SNPs, including information on position, chromosome, allele frequencies, closest gene, type of variant, position relative to closest gene model, if predicted to functionally consequential, and tissues specific gene expression.

### Mouse RBC storage and post-transfusion recovery

Mouse storage studies were performed as previously described ^85^. Initially the analyses were performed on a subset of 350 J:DO mice, then independently repeated by re-analyzing the same 350 along with an additional set of 175 mice to strengthen the analysis, validate and expand the original mQTL findings. All animals were genotyped at 143,259 SNPs (137,192 loci after filtering) using the GigaMUGA array ^104^.

### mQTL analysis in J:DO mice

The mQTL workflow in JAX DO mice followed previously defined conventions ^105^. Briefly, metabolite values of zero were converted to a missing value and only metabolites with 100 non-missing observations or more were included for further analyses. Each metabolite was transformed to normal quantiles for mQTL analysis to reduce the influence of outlying values. The initial mQTL mapping was based on founder strain haplotypes imputed at SNPs (137,192 loci), allowing for the additive genetic effects at loci to be more richly characterized as eight founder allele effects. Sex (43% females and 57% males) and sub-cohorts (5 groups/batches) were adjusted for as covariates. A lenient significance threshold of LOD score > 6 was used for calling mQTL to allow for the detection of mQTL hot spots. For reference, a LOD score > 8 approximately represents a genome-wide adjusted p-value < 0.05 ^106^. Fine mapping at detected mQTL was performed by imputing variants based on the genotypes of the founder strains (GRCm39). We used the same mQTL workflow for fresh and stored RBCs. All analyses were performed using the R package qtl2 ^105^. Processed data and results are uploaded to the QTLViewer webtool (https://churchilllab.jax.org/qtlviewer/Zimring/RBC;^74^) for interactive analyses and download. All data and code for the analysis pipeline are available at figshare (https://doi.org/10.6084/m9.figshare.24456619).

### Dynamic modeling of PFKP

To understand the role of the PFKP isozyme in human RBCs, we used the MASSpy python package to reconstruct a mass action stoichiometric simulation model of the *in vivo* RBC glycolytic pathway at steady state^49,50^. First, we replaced mass action rate laws with detailed enzyme kinetic equations and parameters for key regulatory enzymes hexokinase, phosphofructokinase, and pyruvate kinase^23,53^. Then, we assumed that the existing PFK reaction was catalyzed by the tetrameric enzyme formed by liver and muscle isozymes in erythrocytes. Mature RBCs do not contain significant concentrations of fructose 2,6-bisphosphate, therefore we assumed that fructose 2,6-bisphosphate is negligible as an effector in the intracellular RBC environment. We then created a second copy of the model and modified it with a second PFK reaction to represent the platelet isozyme, whose concentration estimates as a fraction of total PFK isoforms were based on proteomics measurements in the present study. We parameterized the PFKP isozyme in the model relative to the PFKL isozyme from true comparative and protein structure studies where possible to avoid discrepancies due to different assay conditions across studies^51,56–58^. Models representing RBC glycolysis with PFKP and without PFKP were simulated after a sudden decrease in ATP concentration. Extended details and equations in **Supplementary Materials and Methods.**

### Determination of hemoglobin and bilirubin increment via the vein-to-vein database

Association of metabolite levels with hemoglobin and bilirubin increment was performed by interrogating the vein-to-vein database, as described in Roubinian et al. ^30,72^ Bucketing of metabolite levels in quartiles and tertiles for hemoglobin and bilirubin increments was informed by the significantly smaller record of entries for the latter parameter in the database.

### Transfusion Exposures and Outcome Measures

All single RBC unit transfusion episodes linked to genetic polymorphism and metabolomic data were included in this analysis. A RBC unit transfusion episode was defined as any single RBC transfusion from a single donor with both informative pre– and post-transfusion laboratory measures and without any other RBC units transfused in the following 24-hour time period. The outcome measures of interest were change in hemoglobin (ΔHb; g/dL) and change in total bilirubin (ΔBi; mg/dL) following a single RBC unit transfusion episode. These measurements were adjusted for all of the other co-factors – including donor age, sex, hemoglobin – component apheresis, irradiation, storage age, storage solution, recipient age, sex, body mass index, and hemoglobin level. These outcomes were defined as the difference between the post-transfusion and pre-transfusion levels. Pre-transfusion thresholds for these measures were chosen to limit patient confounding (e.g., underlying hepatic disease). For pre-transfusion hemoglobin, the value used was the most proximal hemoglobin measurements prior to RBC transfusion, but at most 24 hours prior to transfusion. Furthermore, we excluded transfusion episodes where the pre-transfusion hemoglobin was greater than 9.5 g/dL, and the hemoglobin increment may be confounded by hemorrhage events. For post-transfusion hemoglobin, the laboratory measure nearest to 24-hours post-transfusion, but between 12– and 36-hours following transfusion was used. For pre-transfusion bilirubin, the most proximal total bilirubin measurement prior to transfusion, but at most 7-days prior to transfusion was used. Furthermore, the pre-transfusion bilirubin had to be below 1.5 mg/dL for the transfusion to be considered to exclude patients with pre-existing liver disease. For post-transfusion bilirubin, the most proximal laboratory measure to the transfusion, but at most 24-hours post-transfusion was used.

### Construction of the glycolytic pathway model

Kinetic models provide an excellent platform for reconciling multiple experimental data types to elucidate underlying mechanisms and relationships within a biologically relevant context. However, the need to reconcile in vitro parameters obtained across different conditions for the parameterization of models creates a series of challenges and limitations associated with the model development, necessitating the tradeoff between model simplicity and predictive power^54^. The following sections describe the construction of the model for exploration of the PFK-P single nucleotide polymorphism within the glycolytic pathway (overview below).

### Glycolytic pathway model for exploring PFKP SNPs (Supplementary Figure 7.G)

The glycolytic pathway model contains 20 metabolites and 23 reactions^107^. Glucose flows into the model while pyruvate, lactate, and inorganic phosphate are exchanged with the environment. The connection to the adenylate salvage pathway is represented by a fixed influx of AMP representing synthesis and a variable efflux representing the corresponding degradation. The load on NADH due to the reduction of methemoglobin are also included. Created using the Escher Network Visualization tool^108^.

We constructed a mass action stoichiometric simulation model of the in vivo RBC glycolytic pathway at steady state using the MASSpy software package^49^. In order to be able to differentiate between isozymes in the model; we replaced the mass action rate law for PFK with a mechanistic expression that explicitly incorporated critical regulatory mechanisms^23^. Previous simulation studies have highlighted the interplay between all three regulatory kinases in rejecting disturbances of glycolytic energy metabolism^109^; we therefore included mechanistic expressions for hexokinase and pyruvate kinase that also incorporated their regulatory mechanisms^23,110,111^.

Next, we added a new reaction to the model to represent the platelet isoform of PFK. PFK is considered to be one of the most important regulatory enzymes of glycolysis, utilizing ATP as both a substrate and an allosteric inhibitor. Each PFK isozyme exhibits differing affinities for substrates and effectors; the simultaneous expression of the different isozymes can therefore expand the cell’s metabolic capacity to respond to various conditions^51,112^. We parameterized the PFK-P isozyme in the model relative to the PFK-L isozyme from true comparative and protein structure studies where possible to avoid discrepancies due to different assay conditions across studies^51,56{Kloos 2015, #644,58^. Mature red blood cells do not contain significant concentrations of fructose 2,6-bisphosphate; therefore we assumed that fructose 2,6-bisphosphate is negligible as an effector in the intracellular RBC environment. ***Assumptions and limitations*** RBCs must recover their ATP content to restore their biophysical properties (e.g., stiffness^59^). We reasoned that if increased PFKP boosted ATP levels at the end of storage, PFKP likely has a role in the initial recovery of cellular ATP. Our study is one of the first to validate and model the presence of PFKP in mature erythrocytes. Once the role of the adenylate pool in the simulated dynamic responses was established, we chose to focus our main analyses on a simpler case study. Through a simpler case study in which the total adenylate pool remains unchanged (**Supplementary Figure 7.H**), the role of PFKP in the observed dynamics is not obfuscated by multiple confounding variables. Our analyses are intended to distinguish the contribution of PFKP relative to PFKL/M, and to provide insight into the significance of the PFKP rs2388595 SNP with respect to the increased ATP levels at the end of storage.

PFK-P was often considered a contaminant in early studies of PFK, there are no existing kinetic studies of PFKP within the context of erythrocytes that we are aware of. Therefore, we chose to parameterize PFKP relative to the erythrocyte PFK. There are also a multitude of technical limitations that hamper the ability to obtain accurate kinetic parameters for the reverse kinetics of PFKP; thus, we assumed the reverse kinetics of PFKP to be negligible, justified by its intrinsic low activity in the forward direction and the large energy barrier that must be overcome for the reverse direction^51,113^.

Our proteomic analyses demonstrated that liver and muscle isozymes comprise ∼89% of the total PFK concentration while the platelet form accounts for the remaining ∼11% (**Figure 6.B**). Therefore, we approximated the total PFKP concentration as 13.6 nM by assuming the measured value of 110 nM ^23^ for erythrocyte PFK represents 89% of the total PFK concentration in the model. In the model without PFKP, the 110 nM value measured for erythrocyte PFK represents 100% of total PFK concentration. The use of simplifying assumptions for parameter reconciliation necessitates caution when interpreting results numerically. However, the observed dynamic responses are consistent with the multi-stable glycolytic behavior and regulatory properties observed in previous *in silico* and experimental studies, providing additional validity to the mechanistic insights gained from simulation.

Modeling isozymes within a cellular context often comes with various difficulties and limitations due to the inability to ascertain the relative load of each isozyme. Isozymes may also differ in catalytic and regulatory properties, further complicated by heterogenous subunit oligomerization, illustrated with the PFK platelet, muscle, and liver subunits^55,58^. As these problems specific to isozymes are in addition to the previously stated need to reconcile in vitro parameters obtained across different conditions, simplifying assumptions become necessary and results must be interpreted within the context of those assumptions.

### Equations and parameters HEX1

Hexokinase (HEX1) is the first key regulatory kinase in glycolysis. The rate equation was taken from^23^. HEX1 is modeled using a partial rapid equilibrium random bi bi mechanism in which all steps in the mechanism equilibrate except the reaction-ternary complexes. Mixed inhibition by 2,3-BPG, GSH, G1,6-BP, and G6P with respect to GLC are also included in the rate equation. The rate equation can therefore be represented as the following:

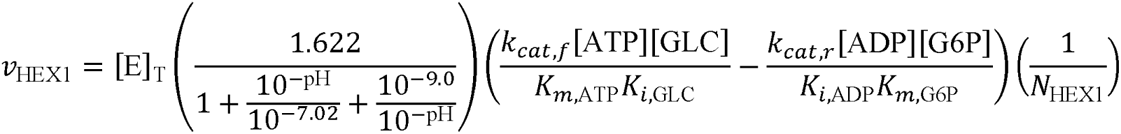

with regulatory component N_HEX1_:

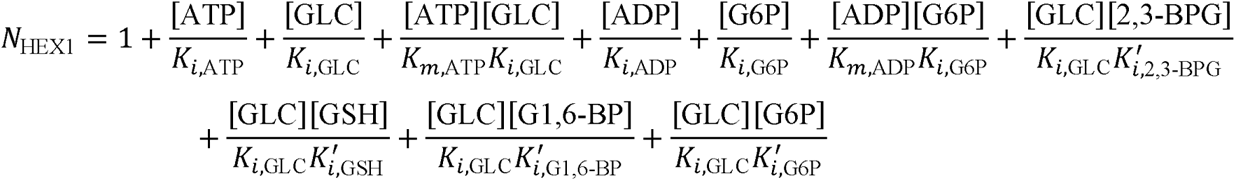

Parameter values were taken from ^23^ unless otherwise stated.

| Parameter | Value |
| --- | --- |
| $K_{m,ATP} = K_{i,ATP}$ | 1 mM |
| $K_{m,GLC} = K_{i,GLC}$ | 0.047 mM |
| $K_{m,ADP} = K_{i,ADP}$ | 1 mM |
| $K_{m,G6P} = K_{i,G6P}$ | 0.047 mM |
| $K'_{i,2,3-BPG}$ | 2.7 mM |
| $K'_{i,GSH}$ | 3.0 mM |
| $K'_{i,G1,6-BP}$ | 0.01 mM |
| $K'_{i,G6P}$ | 0.01 mM |
| $[E]_T$ | $2.5 \times 10^{-5}$ mM |
| $k_{cat,f}$ | $6.48 \times 10^5$ hr<sup>-1</sup> |
| $k_{cat,r}$ | $4.176 \times 10^3$ hr<sup>-1</sup> |
| $K_{HEX1}$ | 850 (Palsson 2011) |
| $k_{PERC, HEX1}$ | $0.7 \text{ mM}^{-1} \text{ hr}^{-1}$ (Palsson 2011) |

### PFK Liver/Muscle

Phosphofructokinase (PFK) is the second key regulatory kinase in glycolysis. The rate equation was taken from^23^. Erythrocyte PFK is modeled using a two-state concerted symmetry mechanism. Inhibition by free ATP, magnesium ion, and 2,3-BPG, and activation by F6P, F1,6-BP, AMP, Pi, and G1,6-BP are also included in the rate equation. Mature RBCs do not contain significant concentrations of fructose 2,6-bisphosphate ^22,27,42,114^; therefore, we assumed that fructose 2,6-bisphosphate is negligible as an effector in the intracellular RBC environment. The explicit complexing of magnesium and free ATP was not included in the model; therefore, we increased the binding constant by a factor of 10, reflecting the ratio of free ATP to MgATP ^107^. The rate equation can therefore be represented as the following:

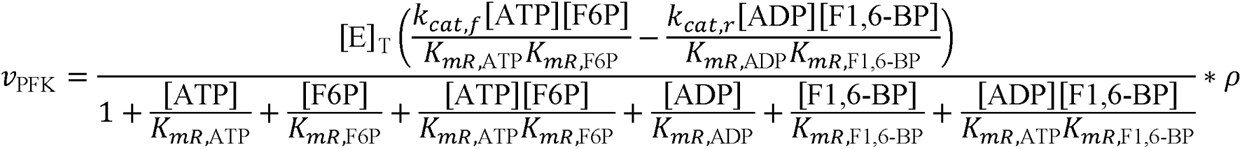

with the active fraction of enzyme *p*:

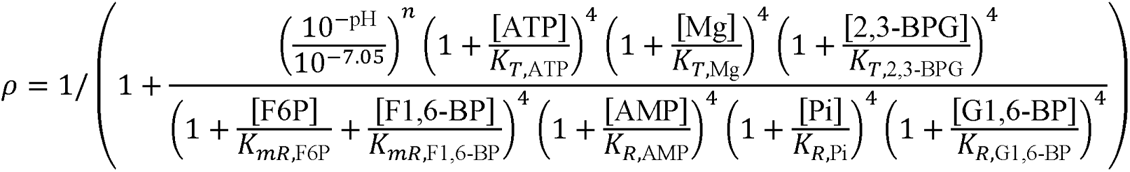

where *R* and *T* represent the relaxed and tense states, respectively. Parameter values were taken from ^23^ unless otherwise stated.

| Parameter | Value |
| --- | --- |
| $K_{mR,ATP}$ | 0.068 mM |
| $K_{mR,F6P}$ | 0.27 mM |
| $K_{mR,ADP}$ | 0.54 mM |
| $K_{mR,F1,6-BP}$ | 0.42 mM |
| $K_{R,AMP}$ | 0.035 mM |
| $K_{R,Pi}$ | 0.431 mM |
| $K_{R,G1,6-BP}$ | 0.0151 mM |
| $K_{T,ATP}$ | 0.098 mM |
| $K_{T,Mg}$ | 0.44 mM |
| $K_{T,2,3-BPG}$ | 1.44 mM |
| $K_a$ | $8.91 * 10^{-8}$ |
| $n$ | 2 |
| $[E]_T$ | $1.1 * 10^{-4}$ mM |
| $k_{cat,f}$ | $2.9592 * 10^6$ hr<sup>-1</sup> |
| $k_{cat,r}$ | $1.296 * 10^5$ hr<sup>-1</sup> |
| $K_{PFK}$ | 310 (Palsson 2011) |
| $k_{PERC, PFK}$ | $35.37$ mM<sup>-1</sup>hr<sup>-1</sup> (Palsson 2011) |

### PFK Platelet isozyme

The platelet isozyme of PFK (PFKP) is distinct among the PFK isozymes as it is not allosterically activated by either fructose 1,6-bisphosphate or glucose 1,6-bisphosphate ^58^. Platelet PFKP is modeled using a modified two-state concerted symmetry mechanism that does not account for the reverse reaction component, as binding affinity for ADP is technically challenging to measure and PFK-P is much less active in the reverse direction ^51,113^. Inhibition by ATP, magnesium ion, and 2,3-BPG, and activation by F6P, AMP, and Pi are also included in the rate equation. Mature RBCs do not contain significant concentrations of fructose 2,6-bisphosphate ^19,22,27,42^; therefore, we assumed that fructose 2,6-bisphosphate is negligible as an effector in the intracellular RBC environment. The rate equation can therefore be represented as the following:

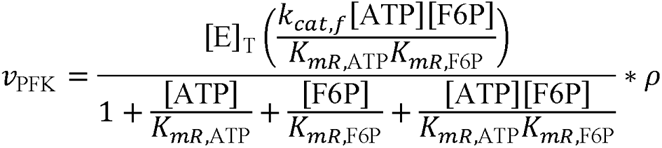

with the active fraction of enzyme *p*:

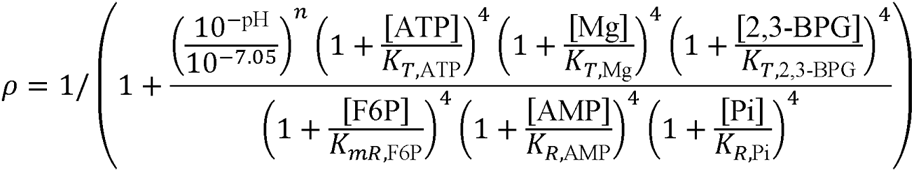

Original parameter values are the same as those for erythrocyte PFK, then modified using the ratio of PFK-L (the most abundant PFK isoform in erythrocytes) to PFK-P values from the structural and comparative studies to reduce differences caused by inconsistencies across experimental conditions. The inhibitory constant for ATP was taken directly from ref.^57^, where it was measured in the presence of magnesium.

| Parameter | Original Value | Modifier<br>(P / L parameter<br>value) | New value | Source |
| --- | --- | --- | --- | --- |
| $K_{mR,\text{ATP}}$ | 0.068 mM | $\frac{275.6 \text{ mM}}{160.1 \text{ mM}}$ | 0.1171 mM | (Fernandes et al. 2020)<sup>51</sup> |
| $K_{mR,\text{F6P}}$ | 0.27 mM | $\frac{1333 \text{ mM}}{1360 \text{ mM}}$ | 0.2645 mM | (Fernandes et al. 2020)<sup>51</sup> |
| $K_{R,\text{AMP}}$ | 0.035 mM | $\frac{0.075 \text{ mM}}{0.010 \text{ mM}}$ | 0.2625 mM | (Foe and Kemp 1985)<sup>115</sup> |
| $K_{T,\text{ATP}}$ | | | 1.6 mM | (Kloos et al. 2015)<sup>57</sup> |
| $[\text{E}]_T$ | $1.1 * 10^{-4} \text{ mM}$ | 0.11 / 0.89 | $1.36 * 10^{-5} \text{ mM}$ | Proteomic analyses,<br>Figure 6.B. |
| $k_{cat,f}$ | $2.9592 * 10^6 \text{ hr}^{-1}$ | $\frac{39.7 * 3600 \text{ hr}^{-1}}{57.3 * 3600 \text{ hr}^{-1}}$ | $2.05027 * 10^6 \text{ hr}^{-1}$ | (Fernandes et al. 2020)<sup>51</sup> |

**PYK** Pyruvate kinase (PYK) is the third key regulatory kinase in glycolysis. The rate equation was taken from ^23^. PYK is modeled using a two-state concerted symmetry mechanism. Inhibition by free ATP and activation by PEP, PYR, F1,6-BP, and G1,6-BP are also included in the rate equation. The explicit complexing of magnesium and free ATP was not included in the model; therefore, we increased the binding constant by factor of 10, reflecting the ratio of free ATP to MgATP ^107^. The rate equation can therefore be represented as the following:

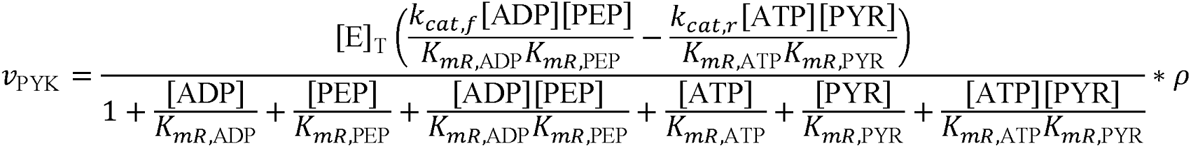

with the active fraction of enzyme *p*:

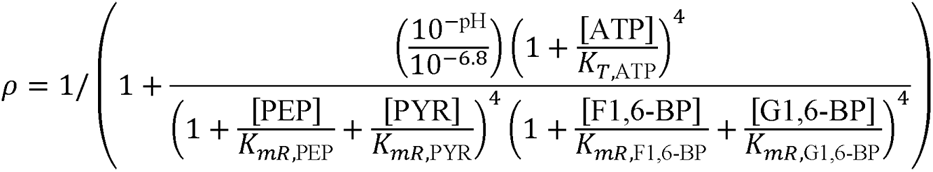

where *R* and *T* represent the relaxed and tense states, respectively. Parameter values were taken from ^23^ unless otherwise stated.

| Parameter | Value |
| --- | --- |
| $K_{mR,\text{ADP}}$ | 0.474 mM |
| $K_{mR,\text{PEP}}$ | 0.225 mM |
| $K_{mR,\text{ATP}}$ | 3.0 mM |
| $K_{mR,\text{PYR}}$ | 2 mM |
| $K_{mR,\text{F1,6-BP}}$ | 0.005 mM |
| $K_{mR,\text{G1,6-BP}}$ | 0.1 mM |
| $K_{T,\text{ATP}}$ | 3.39 * 10 mM |
| $[\text{E}]_T$ | 8.7 * 10<sup>-5</sup> mM |
| $k_{cat,f}$ | 4.9896 * 10<sup>6</sup> hr<sup>-1</sup> |
| $k_{cat,r}$ | 1.1736 * 10<sup>4</sup> hr<sup>-1</sup> |
| $K_{\text{PYK}}$ | 3.63 * 10<sup>5</sup> (Palsson 2011) |
| $k_{\text{PERC, PYK}}$ | 454.39 mM<sup>-1</sup> hr<sup>-1</sup> (Palsson 2011) |

## PGI, FBA, TPI, GAPD, PGK, PGM, ENO, LDH, ADK1

The remaining enzymes equilibrate rapidly without much deviation from their mass action ratios and therefore are modeled using mass action kinetics^116^. Pseudo-elementary rate constants (*k*_PERC_) were computed based on the steady state of the model. Parameter values are taken from ^107^.

| Abbrev. | Description | Rate equation | Kinetic parameters |
| --- | --- | --- | --- |
| PGI | Glucose-6-phosphate isomerase | $v_{\text{PGI}} = k_{\text{PGI}} \left( [\text{G6P}] - \frac{[\text{F6P}]}{K_{\text{PGI}}} \right)$ | $k_{\text{PGI}} = 3644.444$<br>$K_{\text{PGI}} = 0.410$ |
| FBA | Fructose-bisphosphate aldolase | $v_{\text{FBA}} = k_{\text{FBA}} \left( [\text{F1,6-BP}] - \frac{[\text{DHAP}][\text{G3P}]}{K_{\text{FBA}}} \right)$ | $k_{\text{FBA}} = 2834.568$<br>$K_{\text{FBA}} = 0.082$ |
| TPI | Triose-phosphate isomerase | $v_{\text{TPI}} = k_{\text{TPI}} \left( [\text{DHAP}] - \frac{[\text{G3P}]}{K_{\text{TPI}}} \right)$ | $k_{\text{TPI}} = 34.356$<br>$K_{\text{TPI}} = 0.057$ |
| GAPD | Glyceraldehyde-3-phosphate dehydrogenase | $v_{\text{GAPD}} = k_{\text{GAPD}} \left( [\text{G3P}][\text{NAD}][\text{Pi}] - \frac{[\text{1,3-BPG}][\text{NADH}]}{K_{\text{GAPD}}} \right)$ | $k_{\text{GAPD}} = 3376.749$<br>$K_{\text{GAPD}} = 0.018$ |
| PGK | Phosphoglycerate kinase | $v_{\text{PGK}} = k_{\text{PGK}} \left( [\text{1,3-BPG}][\text{ADP}] - \frac{[\text{3-PGLY}][\text{ATP}]}{K_{\text{PGK}}} \right)$ | $k_{\text{PGK}} = 1273531.270$<br>$K_{\text{PGK}} = 1800$ |
| PGM | Phosphoglycerate mutase | $v_{\text{PGM}} = k_{\text{PGM}} \left( [\text{3-PGLY}] - \frac{[\text{2-PGLY}]}{K_{\text{PGM}}} \right)$ | $k_{\text{PGM}} = 4868.589$<br>$K_{\text{PGM}} = 0.147$ |
| ENO | Enolase | $v_{\text{ENO}} = k_{\text{ENO}} \left( [\text{2-PGLY}] - \frac{[\text{PEP}]}{K_{\text{ENO}}} \right)$ | $k_{\text{ENO}} = 1763.741$<br>$K_{\text{ENO}} = 1.695$ |
| LDH | L-lactate dehydrogenase | $v_{\text{LDH}} = k_{\text{LDH}} \left( [\text{PYR}][\text{NADH}] - \frac{[\text{LAC}][\text{NAD}]}{K_{\text{LDH}}} \right)$ | $k_{\text{LDH}} = 1112.57$<br>$K_{\text{LDH}} = 26300$ |
| ADK1 | Adenylate kinase | $v_{\text{ADK1}} = k_{\text{ADK1}} \left( [\text{ADP}]^2 - \frac{[\text{ATP}][\text{AMP}]}{K_{\text{ADK1}}} \right)$ | $k_{\text{ADK1}} = 1112.57$<br>$K_{\text{ADK1}} = 1.65$ |

**ATPM** The non-glycolytic consumption of ATP was modeled as an irreversible process using mass action kinetics. Parameter values are taken from ^107^.

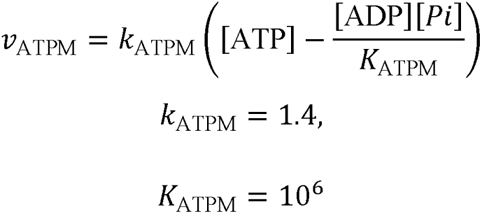

**NADHload** The non-glycolytic consumption of NADH due to the reduction of methemoglobin was modeled as an irreversible process using mass action kinetics. Parameter values are taken from ^107^.

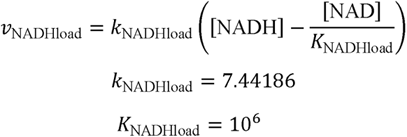

### GLCin, PYRex, LACex, AMPin, AMPout, PIex, Hex, H2Oex

Transport processes in and out of the system were represented using mass action kinetics. Pseudo-elementary rate constants (*k*_PERC_) were computed to reflect a glucose uptake rate of 1.12 mM / hr, an NADH load at 10% of the glycolytic flux (0.224 mM / hr), and an AMP synthesis rate of 0.014 mM /hr. Boundary concentrations are set as constants. Parameter values are taken from ^107^.

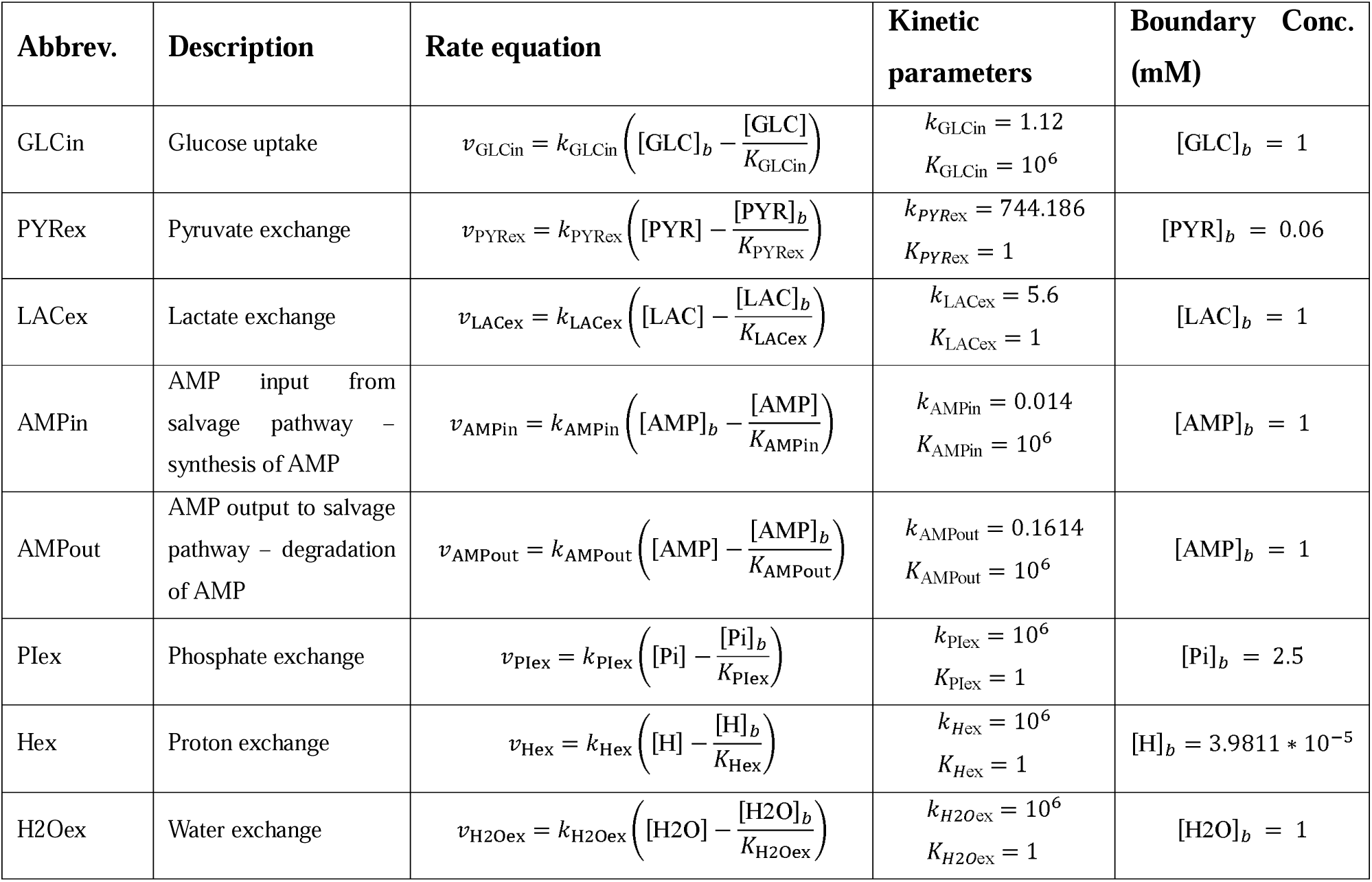

### Description of ordinary differential equations

The ordinary differential equations (ODEs) are derived from the previously defined reaction rate equations. Effector metabolites that are not consumed or produced in the system are held constant at their initial concentration throughout simulation; therefore, their corresponding ODE is set as zero. Initial concentration are taken from ^107^ unless otherwise stated.

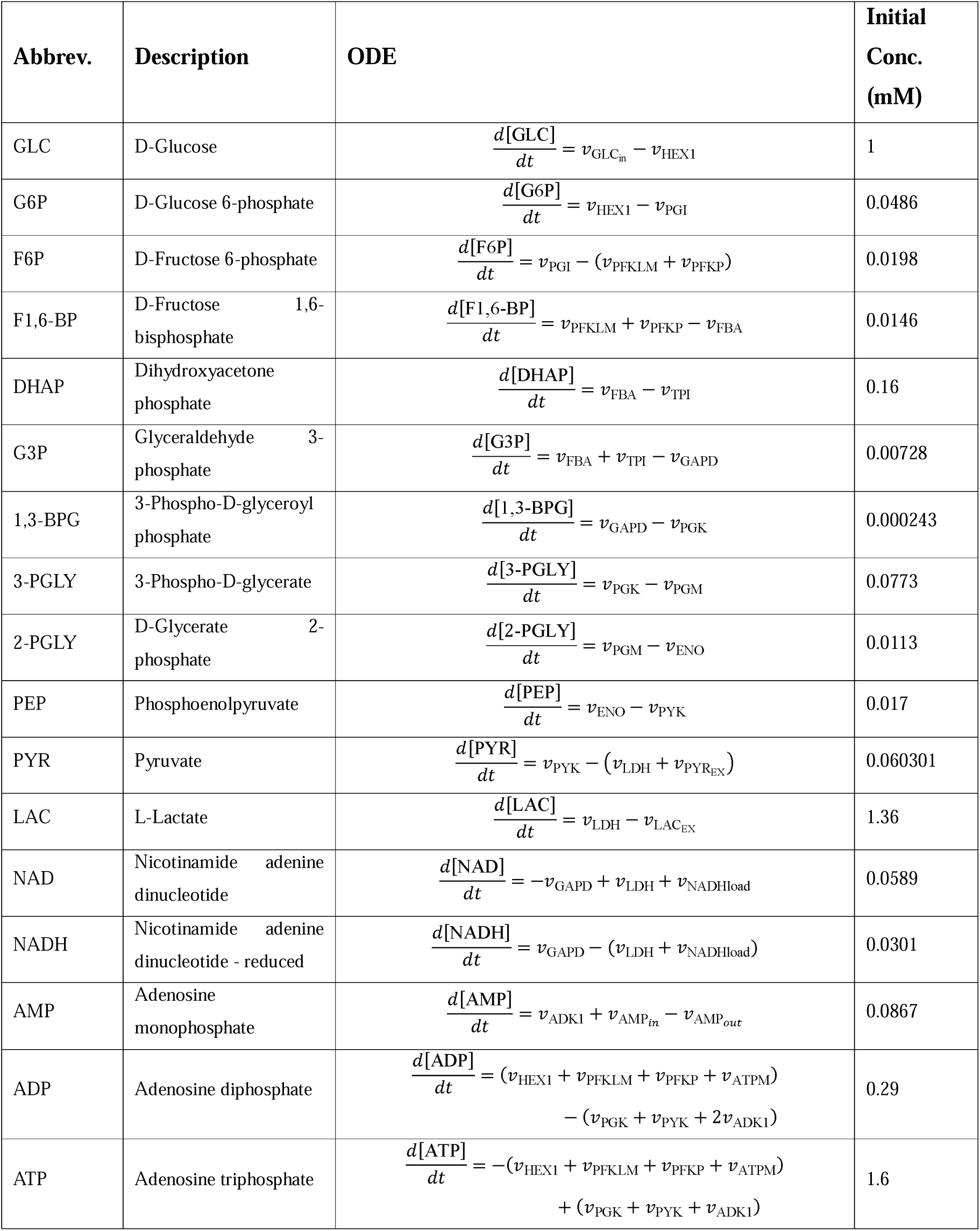
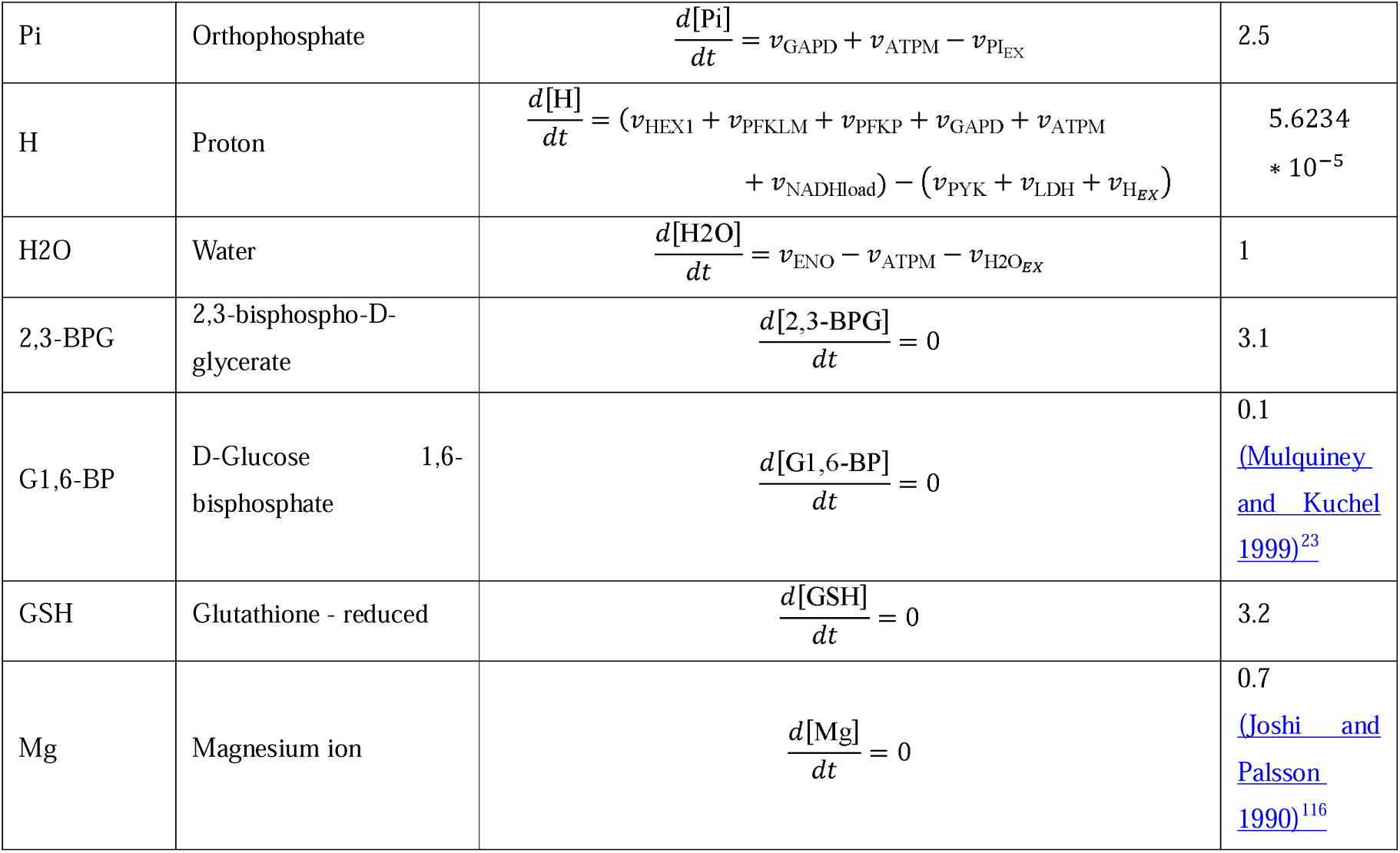

### Adenylate Energy Charge Pools (E.C.)

The adenylate energy charge pools, first described by Atkinson^117^, can be defined as follows (Palsson 2011):

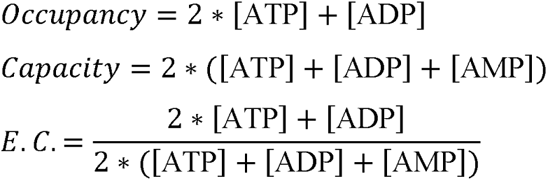

## QUANTIFICATION AND STATISTICAL ANALYSIS

We assessed the univariable association of all *a priori* selected donor, manufacturing, and recipient covariates with the outcomes using linear regression. Multivariable linear regression assessed associations between alleles of donor single nucleotide polymorphisms and changes in hemoglobin and bilirubin levels post-transfusion hemoglobin. Two-sided p-values less than 0.05 were considered to be statistically significant. Analyses were performed using Stata Version 14.1, StataCorp, College Station, TX.

### Data analysis

Statistical analyses – including hierarchical clustering analysis (HCA), linear discriminant analysis (LDA), uniform Manifold Approximation and Projection (uMAP), correlation analyses and Lasso regression were performed using both MetaboAnalyst 5.0^118^ and in-house developed code in R (4.2.3 2023-03-15). Figures were generated in R, Python, MetaboAnalyst, Biorender (www.biorender.com).

