## Supplementary Files for "Biological and Genetic Determinants of Glycolysis: Phosphofructokinase Isoforms Boost Energy Status of Stored Red Blood Cells and Transfusion Outcomes"

**DATA S1-SHEETS 1-11** .......... XLSX
**DATA S1-SHEETS 12-22** .......... XLSX

### SUPPLEMENTARY FIGURES

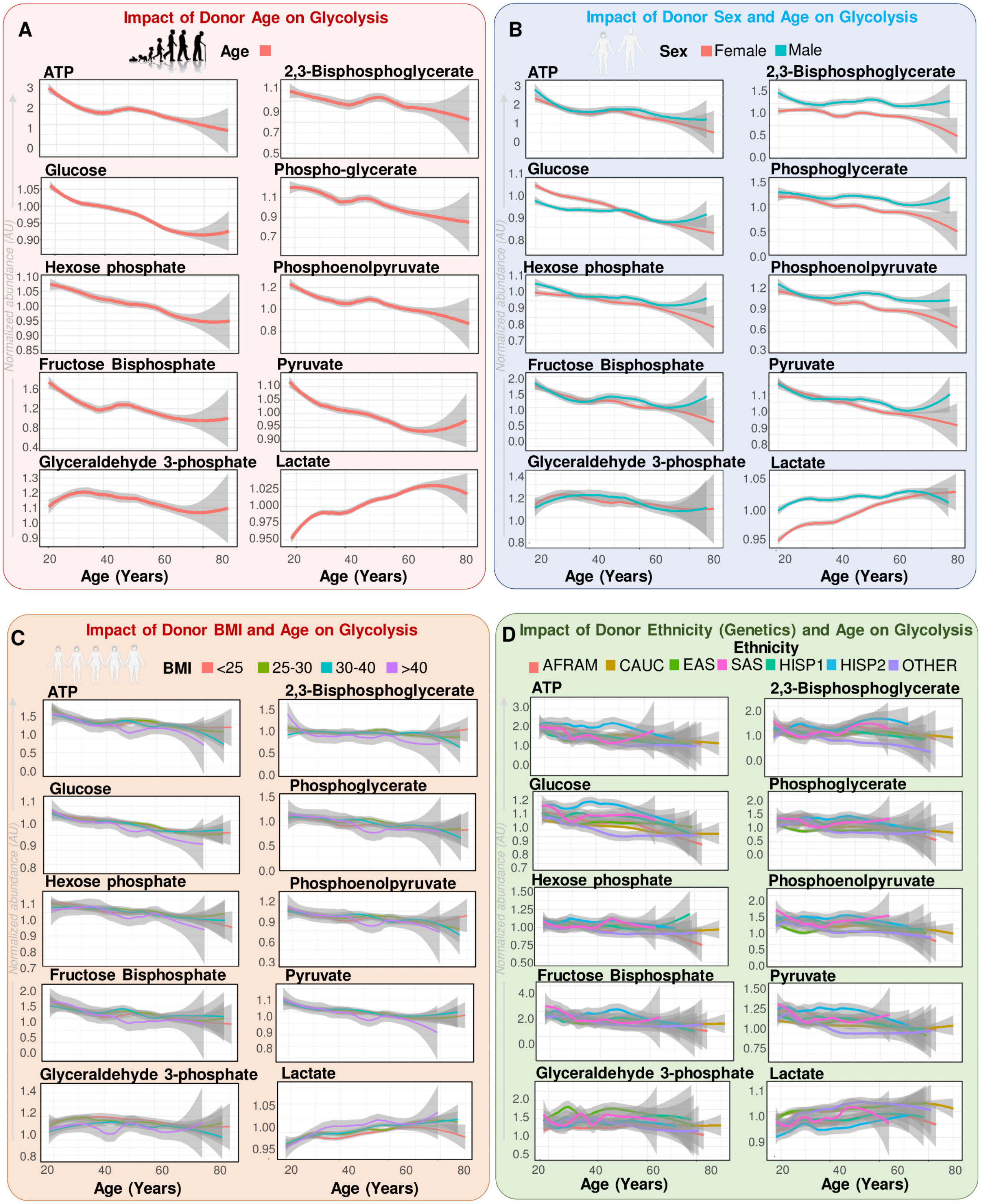

**Supplementary Figure 1 – Glycolysis is impacted by donor sex, age, ethnicity and BMI. Related to Figure 1.** Line plots (median ± quartile range – Loess smoothing algorithm) as a function of donor age, alone (**A**) or in combination with sex (**B**), body mass index (BMI) (**C**) and ethnicity (**D**). Y axes indicate normalized metabolite abundances (arbitrary units – AU). In **D**, abbreviations indicate different ethnicities, including African American (AFRAM); Caucasian (CAUC); Eastern Asian (EAS); South Eastern Asian (SAS); HISP (Hispanic) 1 and 2 – representing Mexican and Central American Hispanics (HISP1) and Caribbean Island Hispanics (HISP2), respectively; and other (mixed ethnic backgrounds).

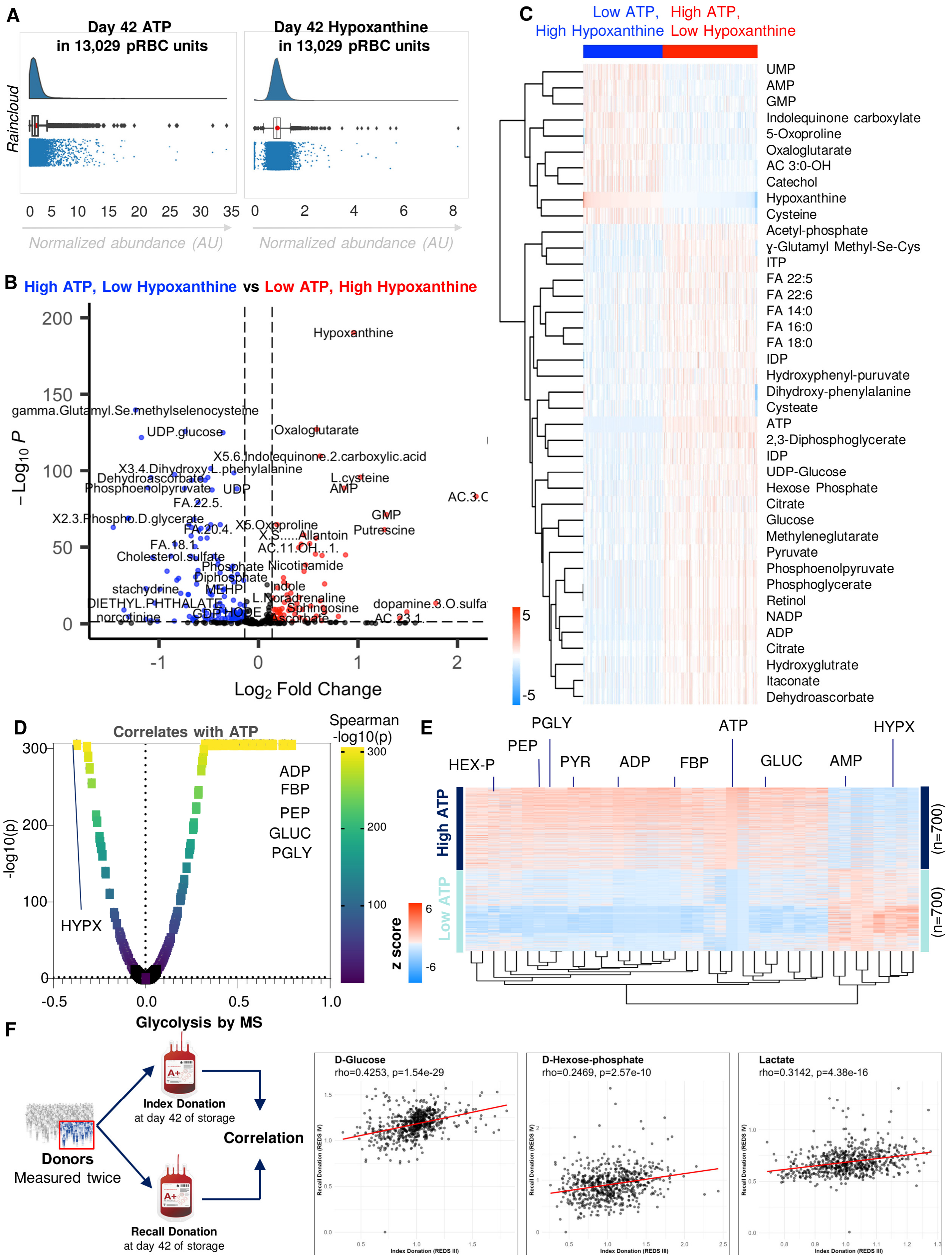

**Supplementary Figure 2** ***Metabolic differences in RBCs from donors with extreme (n=700) highest vs lowest hypoxanthine and concomitantly lowest vs highest ATP.*** **Related to Figure 2.** In **A,** ATP and hypoxanthine distribution in 13,029 pRBC units from the REDS RBC Omics index donors. In **B**, a volcano plot of the differences between the two groups with high vs low hypoxanthine and low vs high ATP. In **C**, heat map of the top 40 metabolites by T-test between the two groups. In **D**, metabolic correlates to ATP levels identify a positive association with glycolytic metabolite and negative with hypoxanthine (HYPX). In **E**, heat map focusing on the ~5% (top 700) donors with highest and lowest hemolysis shows higher levels of glycolytic metabolites and low hypoxanthine proportionally to ATP levels. In **C**, glucose, hexose phosphate and lactate levels are reproducible within the same donor across two independent donations (index vs recalled blood units).

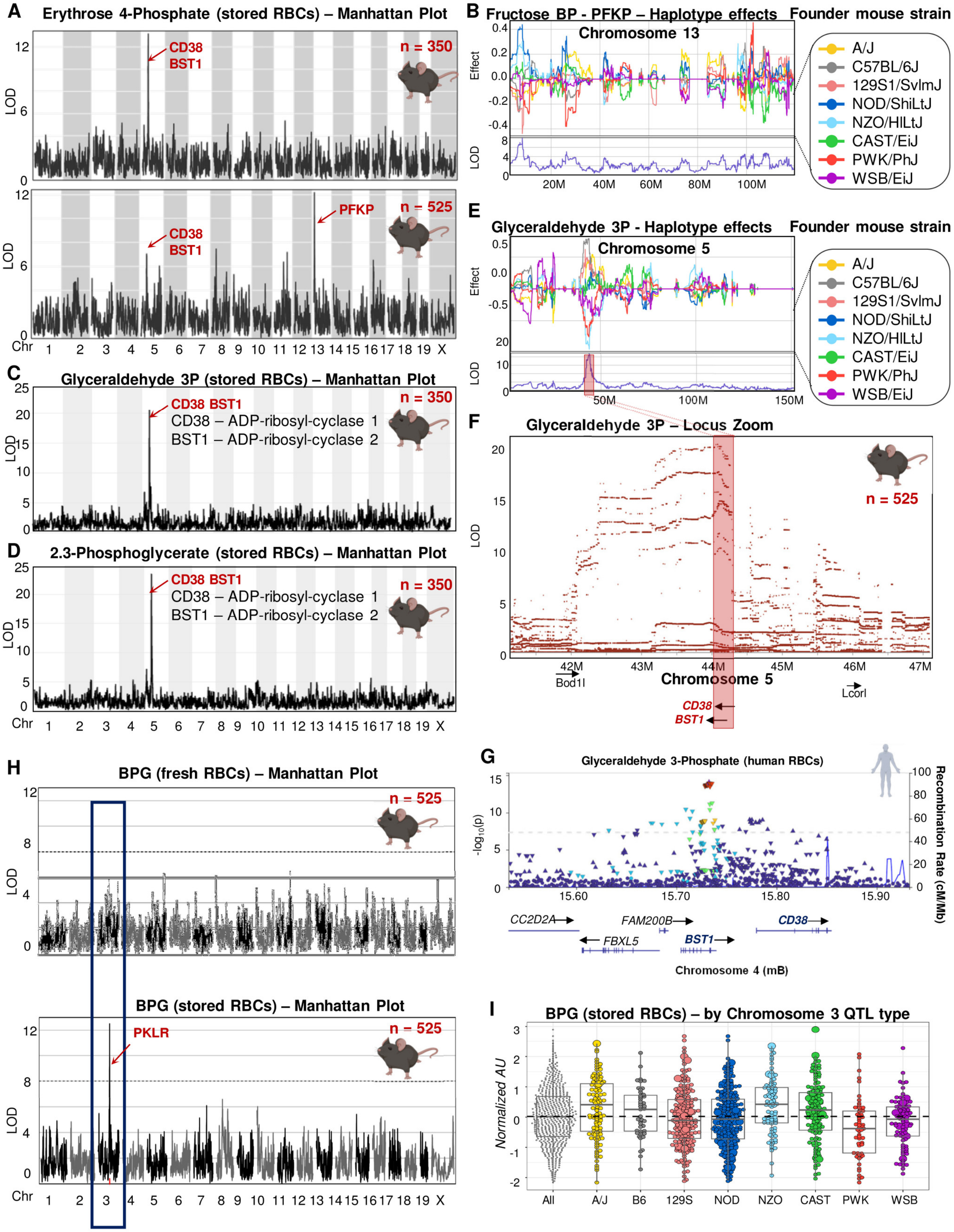

**Supplementary Figure 3** ***CD38/BST1, PFKP and RBC metabolites in stored murine RBCs.*** **Related to Figure 3.**
In **A**, erythrose 4-phosphate association to CD38/BST1 and PFKP was observed in the Jackson laboratory diversity

outbred mice (J:DO) only when testing stored RBC metabolism in 525 mice, but not in a pilot on just 350 mice. In **B**, haplotype effects for fructose bisphosphate (BP) on the region coding for PFKP on murine chromosome 13 based on best linear unbiased predictors (BLUPs). In **C-D**, Manhattan plots for glyceraldehyde 3-phospahte (3P) and 2/3-phosphoglycerate isomers in stored murine RBCs (n=525) identify strong signals on a chromosome 5 region coding for CD38 or BST1 (ADP-ribosyl-cyclase 1 and 2, respectively). Haplotype effects and locus zoom for glyceraldehyde 3-phosphate and chromosome 5 in **E** and **F**. In **G**, locus zoom for glyceraldehyde 3-phosphate in stored human RBCs from Index donors map on a region coding for BST1. In **H**, Manhattan plot for mQTL analyses of BPG in fresh and stored murine RBCs highlights a hit on chromosome 3 in the coding region for pyruvate kinase PKLR, mostly driven by alleles unique to PWK and WSB mice (**I**).

**A**

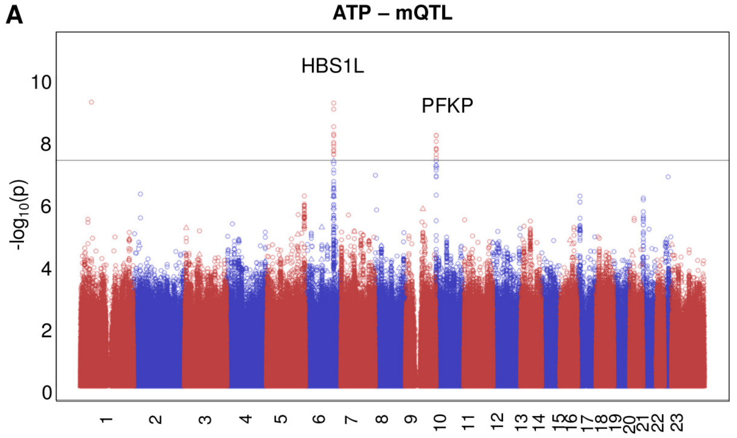

**B**

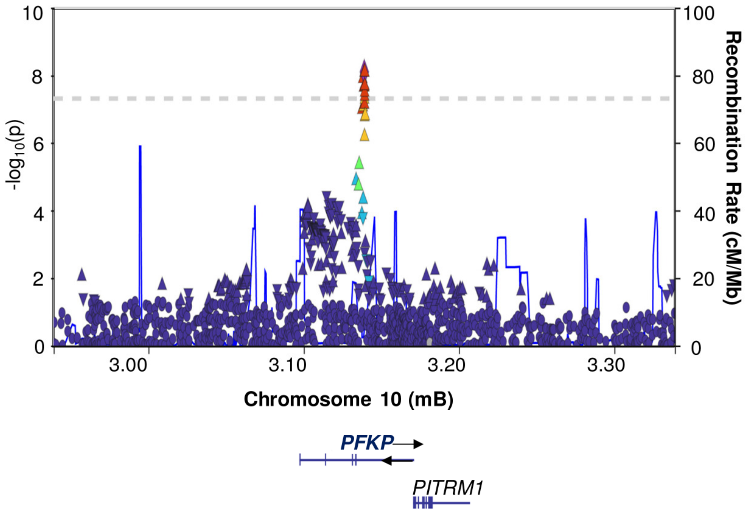

**C**

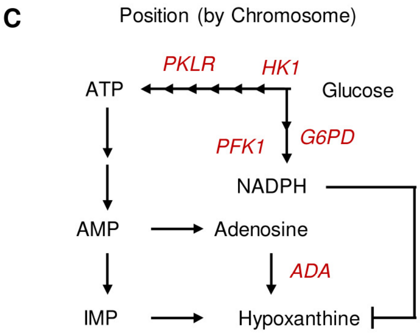

**E**

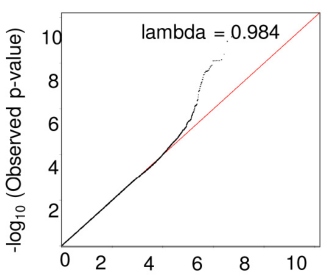

**F** **Minor Allele Frequency for SNP rs2388595**

| | AFR | ASN | EUR | HIS |
|---|---|---|---|---|
| MAF | 0.436 | 0.359 | 0.381 | 0.343 |
| Beta | 0.036499 | 0.016179 | 0.082048 | 0.027287 |
| SE | 0.027315 | 0.028031 | 0.01319 | 0.03616 |
| P | 0.181471 | 0.563825 | 4.96E-10 | 0.450475 |

**D** **Top 10 Significant PFKP SNPs for all glycolytic metabolites combined**

| rsNum | SNP name | p-value | Variant |
|---|---|---|---|
| rs2388595 | 10:3095817:C:T | $1.06 \times 10^{-163}$ | intronic |
| rs2388594 | 10:3095253:G:A | $4.85 \times 10^{-162}$ | intronic |
| rs56882221 | 10:3096285:G:T | $2.97 \times 10^{-161}$ | intronic |
| rs34538474 | 10:3097348:A:G | $7.15 \times 10^{-149}$ | intronic |
| rs10903962 | 10:3096147:A:G | $3.38 \times 10^{-138}$ | intronic |
| rs872398 | 10:3093507:C:T | $7.38 \times 10^{-111}$ | intronic |
| rs7914782 | 10:3098055:A:G | $1.37 \times 10^{-104}$ | intronic |
| rs12249060 | 10:3093094:C:T | $3.94 \times 10^{-95}$ | intronic |
| rs11592619 | 10:3091522:C:T | $4.57 \times 10^{-84}$ | intronic |
| rs55845900 | 10:3100303:T:C | $3.88 \times 10^{-81}$ | intronic |

**G** **Impact of Genetics and Demographics**

Metabolite X ~ age x SNP x sex + HUB + DonFreq

| | Lactate, P | Hypoxanthine, P |
|---|---|---|
| Sex | 2.00E-16 | 0.07 |
| Age | 2.00E-16 | 7.82E-15 |
| SNP | 6.19E-13 | 0.000234 |
| AS-1 | 2.00E-16 | 2.00E-16 |
| AS-3 | 2.00E-16 | 2.00E-16 |
| DonFreq | 1.18E-06 | 1.00E-03 |
| Sex x Age | 1.61E-12 | 4.00E-01 |
| Sex x SNP | 5.00E-01 | 3.00E-01 |
| Age x SNP | 8.00E-01 | 1.00E-01 |

**Supplementary Figure 4 - *mQTL analysis of ATP.* Related to Figure 4.** mQTL analyses were performed for ATP and identified HBS1L (HBS1 like translational GTPase) and PFKP (phosphofructokinase, Platelet) SNPs (**A**). In **B,** locus Zoom for the PFKP coding region. In **C**, schematic summarizing the connection between PFKP, ADA (top hit for hypoxanthine, an ATP breakdown and deamination product – **C**). List of top 10 PFKP SNPs associated with ATP and glycolytic metabolites (especially lactate) by significance (**D**) and related QQ for top PFKP SNPs for ATP and lactate (**E**). In **F**, minor allele frequencies for the rs2388595 PFKP SNP across ancestries. In **G**, mixed mode analysis reporting the effects of donor demographics or storage additives alone, or in combination with PFKP SNP for lactate and hypoxanthine.

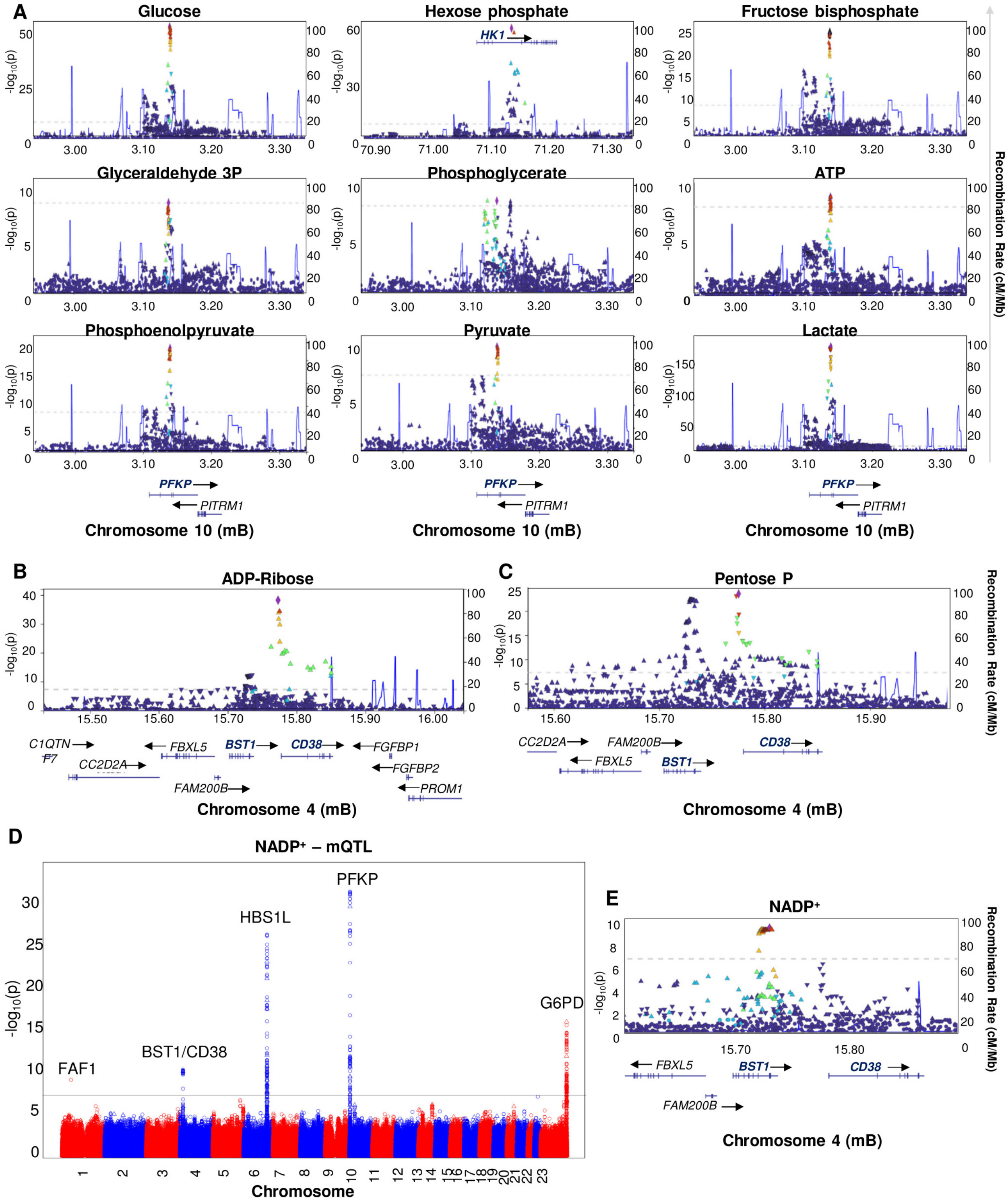

**Supplementary Figure 5** ***Locus Zoom for all the significant SNPs.*** **Related to Figure 4.** for phosphofructokinase, Platelet (PFKP) across all glycolytic metabolites and HK1 (for hexose phosphate only) as determined by mQTL analysis of 13,029 REDS RBC Omics Index donors (**A**). In **B** and **C**, locus zoom plots for ADP-ribose and pentose phosphate showing the region coding for ADP-ribosyl cyclase 1 and 2 (CD38 and BST1, respectively). In **D-E**, Manhattan plot for NADP$^+$ and representative locus zoom for the chromosome 4 region coding for BST1/CD38.

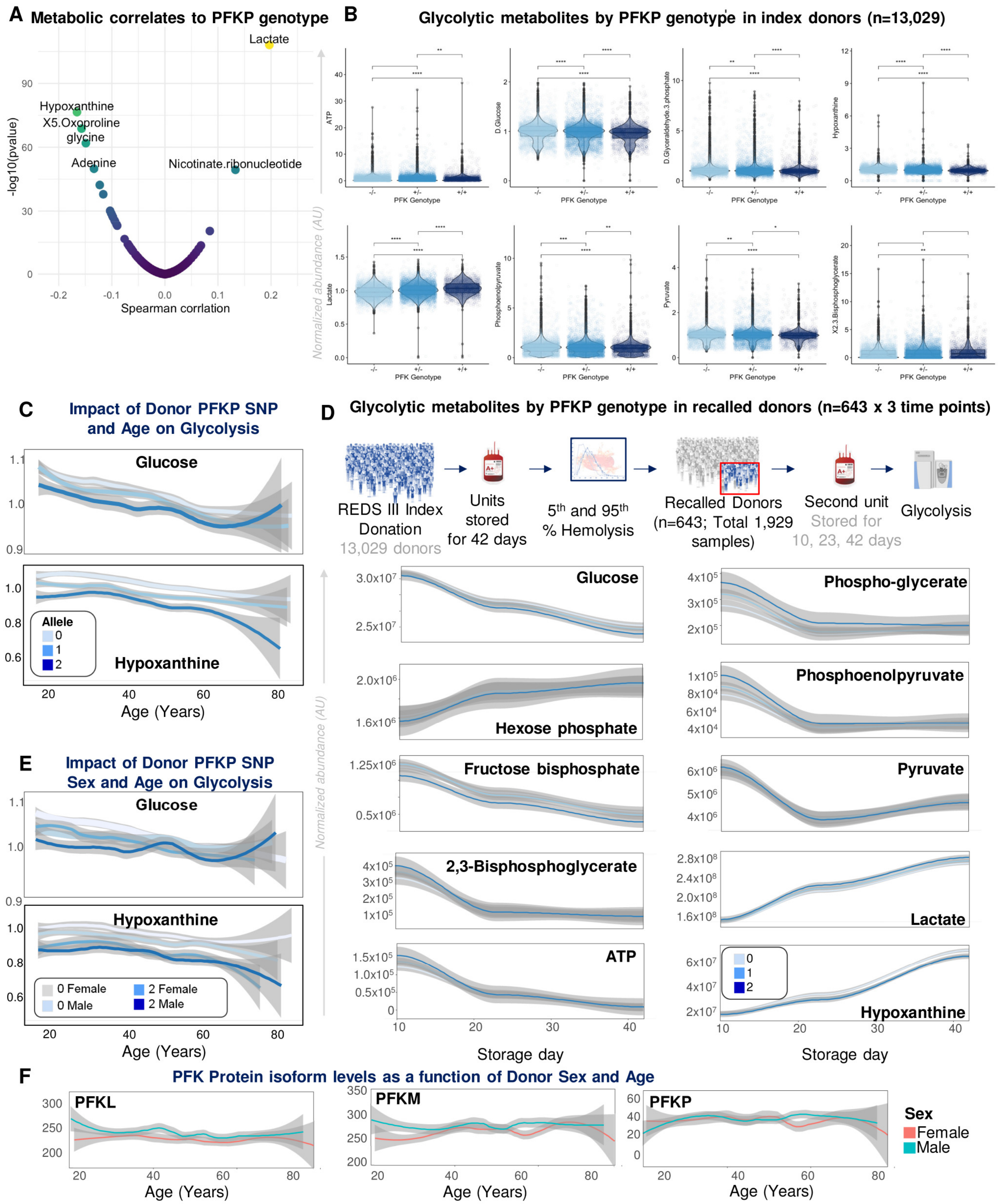

**Supplementary Figure 6 – Metabolic correlates to PFKP SNPs. Related to Figure 6.** PFKP SNP alleles across ~13 thousand REDS index donors were strongly and positively associated with lactate (**A**) and almost all glycolytic metabolites (**B**). Impact of PFKP SNP rs2388595 and donor age alone (**C**) or in combination with sex (**D**) on the end of storage levels of glucose and hypoxanthine in 13,029 blood units from the REDS Index donors. In **E**, glycolytic metabolites at storage day 10, 23 and 42 as a function of PFKP rs2388595 SNP in the REDS Recalled donor population (n=643), as a function of storage age, donor age and sex. No association was observed between PFKP protein expression and donor age or sex in 643 recalled donors (**F**).

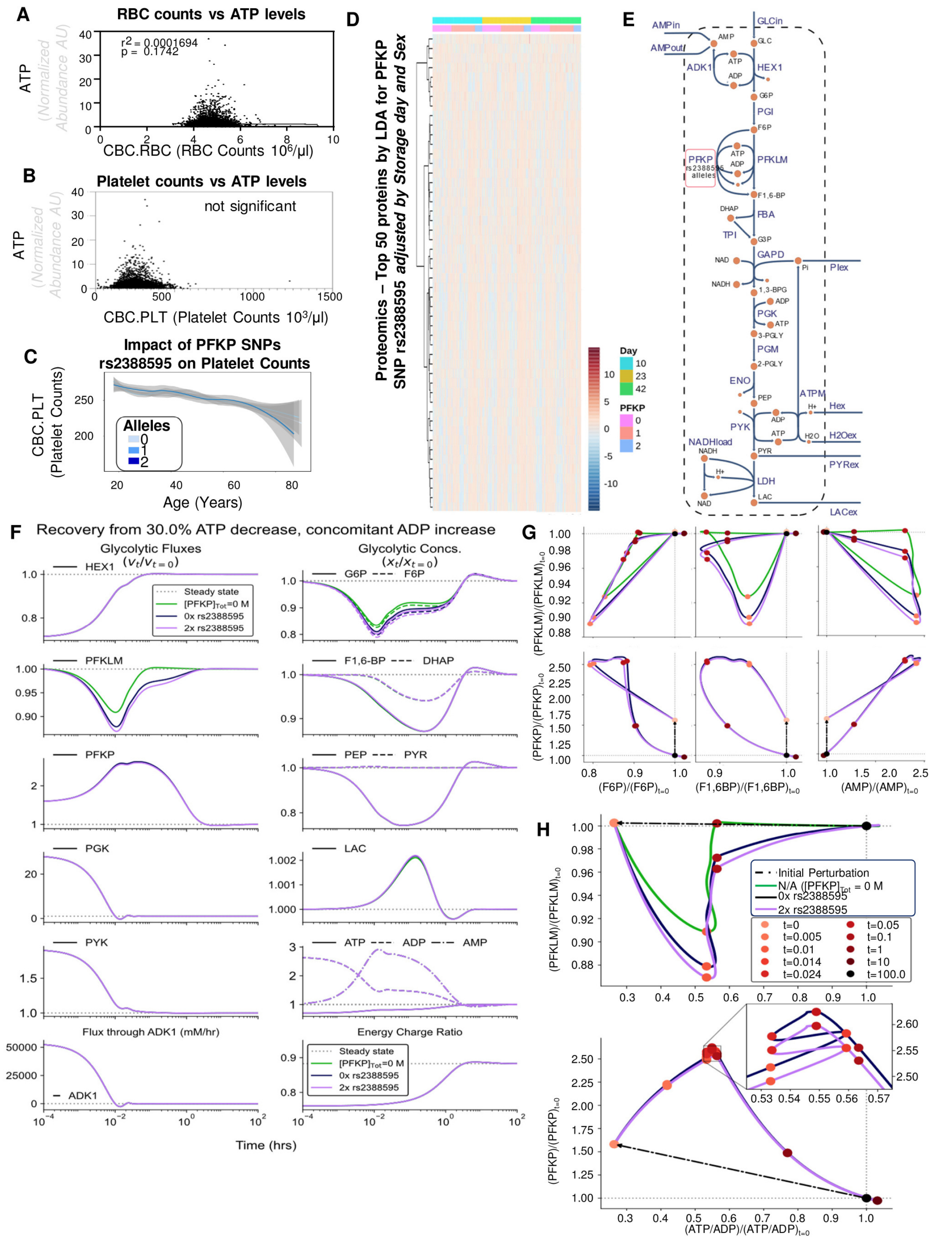

**Supplementary Figure 7 – Metabolic reconstruction of glycolytic fluxes as a function of PFKP genotypes, proteomics and metabolomics data. Related to Figure 6.** ATP levels are not significantly associated with red blood cell (RBC – **A**) or platelet (PLT) counts from cell blood count (CBC) data for REDS RBC Index donors (n=13,029) at the time of donation. The PFKP SNP rs2388595 was not associated with PLT counts in index donors (**C**). No significant changes were observed in the proteome of REDS RBC Omics recalled donors (n=643) during storage (day 10, 23 and 42 – in **D**, top 50 proteins by time-series ANOVA, despite all showing p>0.05). In **G**, glycolytic pathway model for exploring PFKP SNPs. In **E-H**, simulated glycolytic dynamics in the RBC metabolism over time (x axis), with relative contribution of phosphofructokinase (PFK) isoform LM without P (green), and with P in donors carrying 0 (blue) or 2 alleles (purple) of the rs2388595 SNP, based on the PFKP abundance estimates derived from the proteomics data. Results shown for important kinases (hexokinase – HEX1; PFK isoforms LM and P; phosphoglycerate kinase – PGK; pyruvate kinase – PYK; adenylate kinase – ADK1), glycolytic intermediates (glucose 6-phosphate – G6P; fructose 6-phosphate – F6P; fructose 1,6-bisphosphate – F1,6-BP; Dihydroxyacetone phosphate – DHAP; phoshoenolpyruvate – PEP; pyruvate – PYR; lactate - LAC), the adenylate phosphates (ATP, ADP, AMP), and the adenylate energy charge. In **G,** the flux through PFK isoforms LM and P plotted against the F6P, F1,6-BP, and AMP, and (**H**) the ratio of ATP to ADP. Dotted steady state reference lines are included in all plots, with results presented as normalized deviation from steady state unless otherwise noted.
